# IRIS: Big data-informed discovery of cancer immunotherapy targets arising from pre-mRNA alternative splicing

**DOI:** 10.1101/843268

**Authors:** Yang Pan, Alexander H. Lee, Harry T. Yang, Yuanyuan Wang, Yang Xu, Kathryn E. Kadash-Edmondson, John Phillips, Ameya Champhekar, Cristina Puig, Antoni Ribas, Owen N. Witte, Robert M. Prins, Yi Xing

## Abstract

Aberrant alternative splicing (AS) is widespread in cancer, leading to an extensive but largely unexploited repertoire of potential immunotherapy targets. Here we describe IRIS, a computational platform leveraging large-scale cancer and normal transcriptomics data to discover AS-derived tumor antigens for T-cell receptor (TCR) and chimeric antigen receptor T-cell (CAR-T) therapies. Applying IRIS to RNA-Seq data from 22 glioblastomas resected from patients, we identified candidate epitopes and validated their recognition by patient T cells, demonstrating IRIS’s utility for expanding targeted cancer immunotherapy.

## Main

Cancer immunotherapy has gained tremendous momentum in the past decade. The clinical effectiveness of checkpoint inhibitors, such as neutralizing antibodies against PD-1 and CTLA-4, is thought to result from their ability to reactivate tumor-specific T cells^1^. Meanwhile, adoptive cell therapies use genetically modified T-cell receptors (TCRs) or synthetic chimeric antigen receptor T cells (CAR-T) for tumor-specific antigen recognition^2^. The finding that cancer cells express specific T-cell-reactive antigens has galvanized epitope discovery in recent years^3–6^. Nevertheless, the identification of tumor antigens remains a major challenge^7,8^. Although somatic mutation-derived antigens have been successfully targeted by cancer therapies^9–12^, this approach remains largely ineffective for tumors with low or moderate mutation loads^7,13^.

Various types of dysregulation at the RNA level can generate immunogenic peptides in cancer cells^13–15^. Notably, tumors harbor up to 30% more alternative splicing (AS) events than normal tissues, and the resulting peptides are predicted to be presented by human leukocyte antigen (HLA)^16^. However, there are no integrated methods to systematically identify AS-derived tumor antigens. Therefore, we leveraged tens of thousands of normal and tumor transcriptomes generated by large-scale consortium studies (e.g. GTEx, TCGA)^17,18^ to build a versatile, big data-informed platform for discovering AS-derived immunotherapy targets. Our *in silico* platform, named ‘IRIS’ (Isoform peptides from <u>R</u>NA splicing for Immunotherapy target <u>S</u>creening), incorporates three main components: processing of RNA-Seq data, *in silico* screening of tumor AS isoforms, and integrated prediction and prioritization of TCR and CAR-T targets (Fig. 1a).

**Figure 1.**
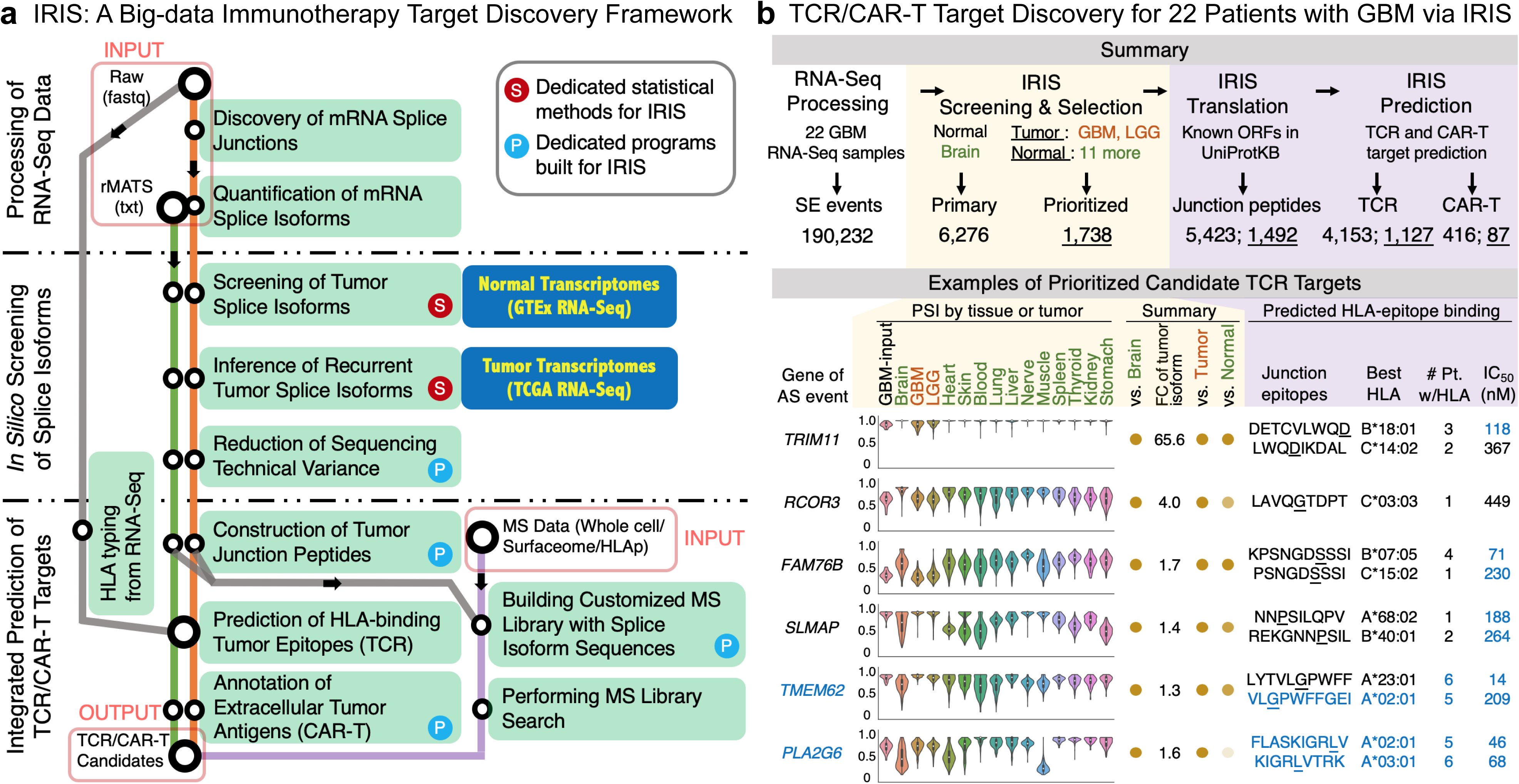
IRIS: A big data-powered platform for discovering AS-derived cancer immunotherapy targets. **a,** Workflow for IRIS, integrating computational modules, large-scale reference RNA-Seq panels, and dedicated statistical testing programs. IRIS has three main modules: RNA-Seq data processing (top), *in silico* screening (middle), and TCR/CAR-T target prediction (bottom). The prediction module includes an option for proteo-transcriptomics integration of RNA-Seq and MS data. **b,** Stepwise results of IRIS to identify AS-derived cancer immunotherapy targets from 22 GBM samples (top). Identified skipped-exon (SE) events from the IRIS data-processing module were screened against tissue-matched normal panel (‘Normal Brain’) to identify tumor-associated events (‘Primary’ set), followed by tumor panel and normal panel to identify tumor-recurrent and tumor-specific events, respectively (‘Prioritized’ set). After constructing splice-junction peptides of tumor isoforms, TCR/CAR-T targets were predicted. As an illustrative example, IRIS readouts for prioritized candidate TCR targets are shown (bottom). Violin plots (left) show PSI values of individual AS events across GBM (‘GBM-input’) versus three reference panels. Dots (middle) summarize screening results. Darker-colored dots indicate stronger tumor features (association/recurrence/specificity) versus each reference panel. FC is estimated fold change of tumor isoform’s proportion in GBM (‘GBM-input’) versus tissue-matched normal panel (‘Brain’). Predicted HLA-epitope binding (right) is output of prediction module. Preferred features for immunotherapy targets in this study are shown in blue. Amino acids at splice junctions in epitopes are underlined. ‘Best HLA’ is HLA type with best predicted affinity (median IC_50_) for given splice-junction epitope. ‘#Pt. w/HLA’ is number of patients with HLA type(s) predicted to bind to a given epitope. Three epitopes in *TMEM62* and *PLA2G6* (blue) were predicted to bind to common HLA types (HLA-A02:01 and HLA-A03:01) and were selected for experimental validation.

IRIS’s RNA-Seq data-processing module uses standard input data to discover and quantify AS events in tumors using our ultra-fast rMATS-turbo software^19,20^. Identified AS events are fed to the *in silico* screening module, which statistically compares AS events against any combination of samples selected from large-scale (>10,000) reference RNA-Seq samples of normal and tumor tissues (Supplementary Fig. 1) to identify AS events that are tumor-associated, tumor-recurrent, and potentially tumor-specific (Methods). Tumor specificity is a key metric for evaluating potential tissue toxicity, which is an important side effect of targeting lineage-specific antigens that are expressed by both tumor and normal cells^21^. In addition to screening multiple patient samples simultaneously in the default ‘group mode’, IRIS can be performed in the ‘personalized mode’ to identify targets for a specific patient sample (Methods). Potential false-positive events are removed by using a blacklist of AS events whose quantification across diverse RNA-Seq datasets is error-prone due to technical variances such as read length (Methods and Supplementary Fig. 2). IRIS’s target prediction module first constructs splice-junction peptides of predicted tumor isoforms and then predicts AS-derived targets for TCR/CAR-T therapies (Methods). This module performs tumor HLA typing using RNA-Seq data and then integrates multiple HLA-binding prediction algorithms for predicting TCR targets and/or peptide vaccines. In parallel, protein extracellular domain annotations are used for predicting CAR-T targets (Supplementary Fig. 3). IRIS also includes the option to confirm predicted AS-derived targets using mass spectrometry (MS) data via proteo-transcriptomics data integration. This option provides an orthogonal approach for target discovery and validation by integrating RNA-Seq data with various types of MS data, such as whole-cell proteomics, surfaceomics, or immunopeptidomics data (Methods and Supplementary Fig. 4a).

We performed a proof-of-concept analysis and preliminary confirmation of AS-derived epitopes by applying IRIS to RNA-Seq and MS-based immunopeptidomics data of cancer and normal cell lines. We identified hundreds of AS-derived epitopes that were supported by both RNA-Seq and MS data (Supplementary Fig. 4b, Supplementary Table 1). MS-supported epitopes were enriched for transcripts with high expression levels and peptides with strong predicted HLA-binding affinities (Supplementary Fig. 4c-e), consistent with the expected pattern of HLA-epitope binding^22^.

To explore IRIS’s ability to discover AS-derived immunotherapy targets in clinical samples, we generated RNA-Seq data from 22 resected glioblastomas (GBMs) and analyzed these data by IRIS. Candidate epitopes were then validated based on their recognition by patient T cells. Fig. 1b (top) summarizes the stepwise IRIS results. After uniform processing of RNA-Seq data by rMATS-turbo, IRIS discovered 190,232 putative skipped exon (SE) events from the 22 GBM samples. Using the *in silico* screening module, we compared these AS events against reference normal and tumor panels to evaluate tumor association, recurrence, and specificity (Methods). Specifically, AS events were compared against: normal brain samples from GTEx (tissue-matched normal panel, for evaluating tumor association), two cohorts of brain tumor samples - GBM and lower-grade glioma (LGG) - from TCGA (tumor panel, for evaluating tumor recurrence), and 11 other selected normal (nonbrain) tissues from GTEx (normal panel, for evaluating tumor specificity). After initially screening against the tissue-matched normal panel and removing blacklisted events, IRIS identified 6,276 tumor-associated AS events in the 22 GBM samples (‘Primary’ set, Fig. 1b). Of these, 1,738 events were identified as tumor-recurrent and tumor-specific based on comparison with the tumor panel and normal panel, respectively (‘Prioritized’ set, Fig. 1b; Supplementary Table 2).

Next, for each AS event, splice junctions of the tumor isoform (i.e. the isoform that was more abundant in the tumor samples than in the tissue-matched normal panel) were translated into peptides, followed by TCR/CAR-T target prediction (Fig. 1b). For the GBM dataset, IRIS predicted 4,153 ‘primary’ tumor-associated epitope-producing splice junctions. Of these, 1,127 were tumor-recurrent and tumor-specific compared to the tumor panel and normal panel, respectively, and were therefore predicted to be ‘prioritized’ TCR targets. In parallel, IRIS identified 416 ‘primary’ tumor-associated extracellular peptide-producing splice junctions, of which 87 were predicted to be ‘prioritized’ CAR-T targets.

IRIS generates an integrative report for predicted immunotherapy targets (Supplementary Table 3). Representative examples for six prioritized TCR targets are shown in the bottom panel of Fig. 1b (see Supplementary Fig. 3b for CAR-T target examples). Violin plots depict exon inclusion levels across the 22 GBM samples (‘GBM-input’) and different sets of reference panels using the percent-spliced-in (PSI) metric^23^. Tumor isoforms can be either the exon-skipped (low PSI) or the exon-included (high PSI) isoform compared to the tissue-matched normal panel. As illustrated by the darker dots in the ‘Summary’ column, all six epitope-producing splice junctions were tumor-associated compared to the tissue-matched normal panel (‘Brain’), and tumor-recurrent compared to the tumor panel (‘GBM’ and ‘LGG’). Two AS events (in *TRIM11* and *FAM76B*) consistently showed distinct PSI values in tumors compared to normal brain and nonbrain tissues, indicating high tumor specificity. For candidate splice junctions, IRIS also calculates the fold-change (FC) of tumor isoforms between tumor samples and the tissue-matched normal panel (Methods). For example, the tumor isoform in *TRIM11* had an average isoform proportion of 8.60% in the 22 GBM samples and 0.13% in normal brain samples, representing an FC of 65.6 in tumor samples versus the tissue-matched normal panel. We should note that, as shown under ‘Predicted HLA-epitope binding’, a single splice junction can give rise to multiple putative epitopes with distinct peptide sequences and HLA binding affinities.

Finally, we sought to validate the immunogenicity and T-cell recognition of IRIS-identified candidate TCR targets using an MHC class I dextramer-based assay^12,24^. We focused on predicted AS-derived tumor epitopes with strong putative HLA-binding affinity to common HLA types found in at least five of the 22 patients. We selected seven AS-derived tumor-associated epitopes (five HLA-A02:01 and two HLA-A03:01) for dextramer-based T-cell recognition testing (Supplementary Table 4). All but one epitope (last one in the table) showed some degree of tumor specificity when evaluated in normal (nonbrain) tissues (‘vs. Normal’, see Fig. 2a). We obtained customized HLA-matched, fluorescently labeled MHC class I dextramer:peptide (pMHC) complexes for each candidate epitope. We conducted flow cytometry to detect CD8^+^ T-cell binding with the pMHC complexes using available peripheral blood mononuclear cells (PBMCs) and/or *ex vivo*-expanded tumor-infiltrating lymphocytes (TILs). Based on the binding of each AS-derived tumor epitope to a patient’s CD3^+^CD8^+^ T cells, we classified epitope reactivity as ‘positive’ (binding > 0.1% of cells), ‘marginal’ (binding 0.01-0.1% of cells), or ‘negative’ (binding < 0.01% of cells). Epitopes that showed at least marginal reactivity were considered to be ‘recognized’ by patient T cells. We analyzed samples from two HLA-A02:01 and four HLA-A03:01 patients, as well as samples from three HLA-A02:01 and three HLA-A03:01 healthy donors (Supplementary Table 5, Supplementary Data).

Both predicted HLA-A03:01 tumor epitopes were recognized by patient T cells. In particular, one epitope (KIGRLVTRK, in *PLA2G6*) was recognized by T cells from all four tested patients but only one of the three tested healthy donors. In one patient (LB2867), recognition of tumor epitope KIGRLVTRK was marginal in PBMCs but positive in the expanded TIL population, with epitope-reactive T cells representing 0.03% of T cells in PBMCs and 1.69% of T cells in TILs. This patient had been previously treated with neoadjuvant anti-PD-1 and anti-CTLA-4 checkpoint blockade immunotherapy. These results suggest epitope KIGRLVTRK as a promising immunotherapy target in HLA-A03 patients from our GBM cohort. T cells from another patient (LB2907) showed positive reactivity to both tested HLA-A03:01 epitopes. All four predicted HLA-A02:01 epitopes were recognized by T cells from tested patients and healthy donors. The non-tumor-specific epitope (YAIVWVNGV, bottom row in Fig. 2a) was tested in two patients and three healthy donors and was recognized by T cells in only one healthy donor (marginal reactivity, 0.013% of CD3^+^CD8^+^ T cells). Taken together, our dextramer-based assay results indicate that the AS-derived TCR targets predicted by IRIS can be recognized by tumor-infiltrating and peripheral CD3^+^CD8^+^ T cells.

Dextramer-positive T cells are expected to contain many clonotypes, only a few of which are dominant. To discover and quantify which TCR clonotypes comprise the epitope-reactive T cells, we sorted the TILs from one patient (LB2867) for cells that reacted positively with the KIGRLVTRK pMHC complex (Fig. 2b), and performed V(D)J immune profiling using single-cell RNA-Seq (scRNA-Seq) on the sorted population (Fig. 2c). Of the 325 unique TCR clonotypes, the 10 most abundant TCRs represented 86.3% of all clonotypes (Supplementary Table 6), with the most frequent clonotype comprising 38.9% of all epitope-reactive T cells. This result suggests that there was clonal expansion of a select few dominant TCR clones within the tumor that were able to recognize the AS-derived epitope. To further validate our findings using complementary approaches, we analyzed bulk expanded TILs using immunoSEQ and pairSEQ assays (Fig. 2c, Supplementary Fig. 5). We confirmed that the top 10 reported clonotypes from scRNA-Seq were present in the bulk TIL population based on the TCR β-chain CDR3 region. In addition, the pairSEQ assay, which uses statistical modeling to predict pairing of TCR α and β chains, found identically paired TCRs for seven of the top 10 TCRs from scRNA-Seq. Together, these data suggest that a select few TCR clones dominantly recognize the AS-derived epitope KIGRLVTRK in this patient.

**Figure 2.**
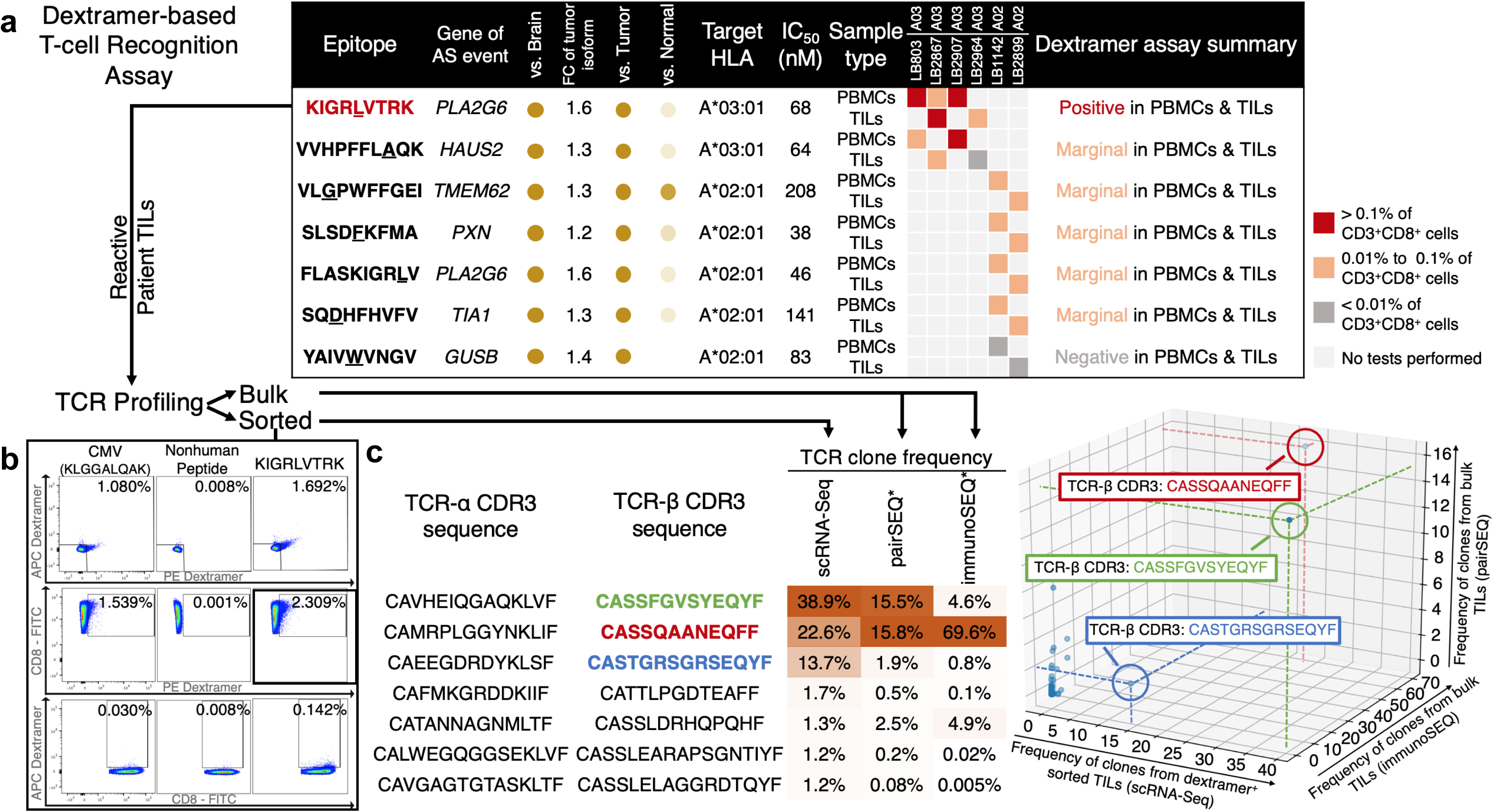
IRIS-predicted AS-derived TCR targets recognized by CD3^+^CD8^+^ T cells in tumors and peripheral blood from patients. **a,** Summary of dextramer-based validation of IRIS-predicted AS-derived epitopes. PBMCs and/or TILs from four HLA-A03 and two HLA-A02 patients were tested for recognition of IRIS-predicted epitopes. Within each HLA type, epitopes are listed by order of tumor specificity (high to low) versus normal panel (11 normal nonbrain tissues). Reactivity (‘Positive’, ‘Marginal’, or ‘Negative’) in assay was evaluated as percentage of dextramer-labeled cells among PBMCs/TILs (>0.1%, 0.01%-0.1%, or <0.01% of CD3^+^CD8^+^ cells, respectively) after subtracting negative control (nonhuman peptide). ‘Dextramer assay summary’ was determined by the mean percent reactivity of CD3^+^CD8^+^ cells across individual tests. **b,** Flow cytometric analysis showing that *ex vivo*-expanded TILs from one HLA-A03 patient (LB2867) contained T cells that recognized epitope KIGRLVTRK. Rows correspond to cells that recognize APC- and PE-labeled dextramers (top), only PE-labeled dextramers (middle), or only APC-labeled dextramers (bottom). Percentages of epitope-specific cells are shown. **c,** Immune profiling results revealing immune repertoire composition of KIGRLVTRK-specific T cells from one patient (LB2867). The scRNA-Seq assay was performed on sorted KIGRLVTRK-specific T cells, whereas pairSEQ and immunoSEQ assays captured TCR clones from bulk TIL RNAs of same patient. Table (left) lists seven most abundant T-cell clones from scRNA-Seq, with percentages of matching CDR3 sequences from TCR β chains. *For pairSEQ and immunoSEQ, percentages are the best frequencies of matching TCR pair or β-chain clones. The 3D scatterplot (right) shows that these approaches converged on three dominant TCR clones. For comparison, the same epitope in the table and 3D scatterplot are identified by use of the same color for the sequence (table) and text box (plot).

In summary, we have developed IRIS, a big data-powered platform for discovering AS-derived tumor antigens as an underexploited source of immunotherapy targets. Using IRIS followed by a dextramer-based assay, we discovered and validated AS-derived tumor epitopes recognized by T cells in patients. Our results provide experimental evidence for the immunogenicity of tumor antigens arising from AS and reveal novel potential targets for TCR and CAR-T therapies. The IRIS software can be downloaded from https://github.com/Xinglab/IRIS.

## Methods

### IRIS module for RNA-Seq data processing

IRIS accepts standard formats of raw RNA-Seq FASTQ files and/or tab-delimited files of quantified AS events (from rMATS-turbo) as input data (Fig. 1a). For raw RNA-Seq data, IRIS provides a standalone pipeline that aligns RNA-Seq reads to the reference human genome hg19 using the STAR 2.5.3a^25^ two-pass mode, followed by Cufflinks v2.2.1^26^ and rMATS v4.0.2 (rMATS-turbo)^19,20^ for quantification of gene expression and AS events, respectively, based on the GENCODE (V26)^27^ gene annotation. To quantify AS events, we converted splice-junction counts in rMATS-turbo output into PSI^23^ values. For each dataset, we removed low-coverage AS events, defined as events with an average count of less than 10 reads for the sum of all splice junctions across all samples in that dataset (tissue/tumor type). We applied this procedure to the 22 GBM samples from the UCLA cohort (BioProject: PRJNA577155), as well as to the normal and tumor samples of the reference panels used by IRIS. For the GTEx normal samples, aligned BAM files downloaded from the dbGAP repository were used directly for AS quantification.

### Constructing big-data reference panels of AS events across normal human tissues and tumor samples

IRIS’s big-data reference panels of normal and tumor samples are available as pre-processed, pre-indexed databases for fast retrieval by the IRIS program (Supplementary Fig. 1). Specifically, 9,662 normal samples from the GTEx project (V7)^17^ representing 53 tissue types of 30 histological sites were uniformly processed as described above. As shown in Supplementary Fig. 1a-b, exon-based quantification of AS events was able to distinguish samples by tissue type. Selected TCGA^16,28^ tumor samples (Supplementary Fig. 1c) were processed similarly to form the tumor panel. Additionally, IRIS provides a stand-alone indexing function for users to include custom normal and tumor samples in their reference panels.

### IRIS module for *in silico* screening of tumor AS events

IRIS performs *in silico* screening using two-sided and one-sided *t*-tests to identify tumor-associated, tumor-recurrent, and tumor-specific AS events in group comparisons. To define an AS event as significantly different from a reference group (i.e., to identify tumor-associated/tumor-specific events), IRIS sets two requirements: 1) a significant p-value from the two-sided *t*-test (default: p < 0.01), and 2) a threshold of PSI value difference (default: abs(ΔΨ) > 0.05). With a minor modification, to define an AS event as tumor-recurrent, IRIS compares a reference tumor panel with the tissue-matched normal panel and requires: 1) a significant p-value from the one-sided *t*-test in the same direction as the corresponding ‘tumor-associated’ event (default: p < 0.01/number of ‘tumor-associated’ events; Bonferroni correction applied due to large sample sizes in reference panels), and 2) a threshold of PSI value difference (default: abs(ΔΨ) > 0.05). In addition, as the normal or tumor reference panel may include multiple individual groups (e.g. tissue types), a threshold of the number of significant comparisons against groups in the normal or tumor reference panel is used to determine whether AS-derived antigens are tumor-specific or tumor-recurrent. For each AS event, IRIS defines the ‘tumor isoform’ as the isoform that is more abundant in tumors than in the tissue-matched normal panel. Optionally, to rank or filter targets, IRIS estimates the ‘fold-change (FC) of tumor isoform’ as the FC of the tumor isoform’s proportion in tumors compared to the tissue-matched normal panel. In addition to the default ‘group mode’, IRIS can be used to screen targets for a specific patient sample through the ‘personalized mode’. This mode uses an outlier detection approach, combining a modified Tukey’s rule^29^ and a user-defined threshold of PSI value difference.

### Identification of AS events that are prone to measurement errors due to technical variances across big-data reference panels

IRIS’s big-data reference panels were constructed by integrating various large-scale datasets with distinct technical conditions, such as RNA-Seq read length^30^. Such technical variances across datasets could introduce discrepancies in the quantification of AS events^30^. To identify error-prone AS events, we employed a data-based heuristic strategy to assess the effects of RNA-Seq read length (48 bp vs. 76 bp) and aligner (STAR vs. Tophat) on AS quantification (PSI value) (Supplementary Fig. 2a). For a given tissue type (in this study, brain tissue), 10 randomly selected 76-bp RNA-Seq files from GTEx were artificially trimmed to 48 bp, and both 76- and 48-bp RNA-Seq files were aligned with STAR2.5.3a. Corresponding Tophat (v.1.4.1)-aligned 76-bp BAM files were directly downloaded from GTEx. AS events were quantified by rMATS-turbo. Events with significantly different PSI values (p < 0.05, abs(ΔΨ) > 0.05 from paired *t*-test) among RNA-Seq datasets with distinct technical conditions were included in a blacklist. Results of this analysis for GTEx normal brain samples are shown in Supplementary Fig. 2b.

### IRIS module for predicting AS-derived TCR and CAR-T targets

To obtain protein sequences of AS-derived tumor isoforms, IRIS generates peptides by translating splice-junction sequences into amino-acid sequences using known ORFs from the UniProtKB^31^ database. Within each AS event, the splice-junction peptide sequence for the tumor isoform is compared to that of the alternative normal isoform, to ensure that the tumor isoform splice junction produces a distinct peptide.

For TCR target prediction, IRIS employs seq2HLA^32^, which uses RNA-Seq data to characterize HLA class I alleles for each tumor sample. IRIS then uses IEDB API^33^ predictors to obtain the putative HLA binding affinities of candidate epitopes. The IEDB ‘recommended’ mode runs several prediction tools to generate multiple predictions of binding affinity, which IRIS summarizes as a median IC_50_ value. By default, a threshold of median(IC_50_) < 500 nM denotes a positive prediction for an AS-derived TCR target.

For CAR-T target prediction, IRIS maps AS-derived tumor isoforms to known protein extracellular domains (ECDs), as potential candidates for CAR-T therapy (Supplementary Fig. 3a). Specifically, IRIS generates pre-computed annotations of protein ECDs. First, protein cellular localization information was retrieved from the UniProtKB^31^ database (flat file downloaded in April 2018). ECD information was retrieved by searching for the term ‘extracellular’ in topological annotation fields, including ‘TOPO_DOM’, ‘TRANSMEM’, and ‘REGION’, in the flat file. Second, BLAST^34^ was used to map individual exons in the gene annotation (GENCODE V26) to proteins with topological annotations. Third, the BLAST result was parsed to create annotations of the mapping between exons and ECDs in proteins. These pre-computed annotations are queried to search for AS-derived peptides that can be mapped to protein ECDs as potential CAR-T targets.

### Proteo-transcriptomics data integration for MS validation

IRIS includes an optional proteo-transcriptomics data integration function that incorporates various types of MS data, such as whole-cell proteomics, surfaceomics, or immunopeptidomics data, to validate RNA-Seq-based target discovery at the protein level (Supplementary Fig. 4a). Specifically, sequences of AS-derived peptides are added to canonical and isoform sequences of the reference human proteome (downloaded from UniProtKB in September 2018). For immunopeptidomics data, fragment MS spectra are searched against the RNA-Seq-based custom proteome library with no enzyme specificity using MSGF+^35^. The search length is limited to 7-15 amino acids. The target-decoy approach is employed to control the false discovery rate (FDR) or ‘QValue’ at 5%.

### IRIS analysis of immunopeptidomics data

Published matching RNA-Seq and MS immunopeptidomics data of B-LCL-S1 and B-LCL-S2 cell lines (B lymphoblastoid cell lines from two individual donors) were retrieved from Laumont *et al.*^36^ (GEO: GSM1641206, GSM1641207, and PRIDE: PXD001898). Raw RNA-Seq data of the JeKo-1 lymphoma cell line were obtained from the Cancer Cell Line Encyclopedia via the NCI Genomic Data Commons (https://portal.gdc.cancer.gov/legacy-archive/). Corresponding immunopeptidomics MS data of JeKo-1 were retrieved from Khodadoust *et al.*^37^ (PRIDE: PXD004746).

RNA-Seq data of the normal (B-LCL-S1, B-LCL-S2) and cancer (JeKo-1) cell lines were analyzed by IRIS as described above, with minor modifications. Specifically, AS events identified by the IRIS RNA-Seq data processing module were not subjected to the *in silico* screening module, but instead were directly used for the MS search. For MSGF+, FDR was set at 5%, which had the best concordance with predicted binding affinities (Supplementary Fig. 4c-d). For comparison of predicted HLA binding and nonbinding peptides (Supplementary Fig. 4d), a set of nonbinding peptides was created by randomly selecting peptides with median(IC_50_) > 500 nM to the same number of binding peptides (median(IC_50_) < 500 nM).

### IRIS discovery of candidate TCR and CAR-T targets from 22 GBM samples

RNA-Seq samples were processed by IRIS. Detected skipped exon (SE) events were analyzed by using the IRIS screening and target prediction modules with the aforementioned default parameters. For reference panels, the ‘tissue-matched normal panel’ comprised normal brain tissue samples from GTEx; the ‘normal panel’ comprised other normal (nonbrain) tissue samples of 11 selected vital tissues (heart, skin, blood, lung, liver, nerve, muscle, spleen, thyroid, kidney and stomach) from GTEx; and the ‘tumor panel’ comprised two cohorts of brain tumor samples (GBM and LGG) from TCGA. The blacklist of AS events created for brain was applied before *in silico* screening by IRIS to eliminate error-prone AS events (Supplementary Fig. 2).

In screening for the ‘Primary’ set of AS events, we considered an event to be ‘tumor-associated’ if it was significantly different from the tissue-matched normal panel, using the default criteria described in ‘IRIS module for *in silico* screening of tumor AS events’. In screening for the ‘Prioritized’ set, we prioritized an AS event if it was both ‘tumor-recurrent’ (significantly different from the tissue-matched normal panel, in the same direction as input GBM samples, in at least 1 of 2 groups in the GBM/LGG tumor panel) and ‘tumor-specific’ (significantly different from multiple of 11 groups in the normal panel in the same direction as the tissue-matched normal panel). Here, we used at least 2 groups but this threshold can be user-defined to allow for higher stringency.

When selecting potential TCR targets for dextramer validation, we applied three additional criteria: 1) predicted median(IC_50_) ≤ 300 nM; 2) predicted binding to common HLA types, including HLA-A02:01 and HLA-A03:01; and 3) predicted binding to at least five patients in the GBM cohort. After excluding targets with low gene expression (average FPKM < 5), we selected seven epitopes to test for T-cell recognition by dextramer assays.

### Patients

Tumor specimens were collected from 22 consenting patients with GBM who underwent surgical resection for tumor removal at the University of California, Los Angeles (UCLA; Los Angeles, CA). From these patients, we also obtained PBMCs and TILs from two HLA-A02:01+ and four HLA-A03:01+ patients. All patients provided written informed consent, and this study was conducted in accordance with established Institutional Review Board-approved protocols.

### PBMC collection

Peripheral blood was drawn from patients before surgery and diluted 1:1 in RPMI media (Thermo Fisher Scientific, cat. no. MT10041CV). PBMCs, extracted by Ficoll gradient (Thermo Fisher Scientific, cat. no. 45-001-750), were washed twice in RPMI media. Collected PBMCs were frozen in 90% human AB serum (Thermo Fisher Scientific, cat. no. MT35060CI) and 10% DMSO (Sigma, cat. no. C6295-50ML) and stored in liquid nitrogen. In parallel, PBMCs from healthy HLA-A02:01 and HLA-A03:01 donors were purchased from Bloodworks Northwest (Seattle, WA) or Astarte Biologics (Bothell, WA).

### TIL collection

Surgically resected tumor samples were digested with a brain tumor dissociation kit (Miltenyi Biotec, cat. no. 130-095-42) and gentle MACS dissociator (cat. no. 130-093-235). After digestion and myelin depletion, collected cells were labeled with CD45 microbeads (cat. no. 130-045-801) and separated on Miltenyi LS columns (cat. no. 130-042-401) and MidiMACS Separator (cat no. 130-042-302). Collected CD45^+^ cells were cultured at 1×10^6^ cells/mL in X-VIVO 15 Media (Fisher Scientific, cat. no. BW04-418Q) containing 2% human AB serum with 50 ng/mL anti-CD3 antibody (BioLegend, cat. no. 317304), 1 μg/mL anti-CD28 antibody (BD Biosciences, cat. no. 555725), 1 μg/mL anti-CD49d antibody (BD Biosciences, cat. no. 555501), 300 IU/mL IL-2 (NIH, cat. no. 11697), and 10 ng/mL IL-15 (BioLegend, cat. no. 570302). Cells were expanded for 3-4 weeks and replenished with fresh media and cytokines every 2-3 days. Before freezing, expanded cells were placed in media containing 50 IU/mL IL-2 for 1-2 days and then frozen in the same freezing media as PBMCs.

### Collection of tumor RNAs and RNA sequencing

RNA from freshly collected or flash-frozen tumor specimens was extracted by using the RNeasy Mini Kit (Qiagen, cat. no. 74014). Paired-end RNA-Seq was performed at the UCLA Clinical Microarray Core using an Illumina HiSeq 3000 at a read length of 2×100 bp or 2×150 bp.

### Dextramer flow-cytometric analysis of PBMCs and TILs

For each AS-derived peptide selected for validation, custom-made HLA-matched MHC Class I dextramer:peptide (pMHC) complexes were purchased from Immudex (Copenhagen, Denmark). Immudex also provided pMHC complexes for common cytomegalovirus (CMV) epitopes (cat. nos. WB2132 and WC2197) and for a nonhuman epitope (NI3233) as a negative control. Each pMHC complex was purchased with two separate tags for APC or PE fluorescence labeling, to increase specificity to targeted T cells with dual labeling.

To facilitate proper gating of CD8^+^ T cells from PBMC and TIL populations, the following panel of antibodies (from BioLegend) was set up: CD3 BV605 (cat. no. 300460), CD8 FITC (cat. no. 344704), CD4 BV421 (cat. no. 317434), CD19 BV421 (cat. no. 302234), CD56 BV421 (cat. no. 362552), and CD14 BV421 (cat. no. 301828). For single-color compensation controls, OneComp eBeads were used (Thermo Fisher Scientific, cat. no. 01-1111-41).

For each set of pMHC complexes, at least 3×10^6^ cells were stained according to manufacturer’s guidelines. Briefly, cells were thawed in a 37°C water bath and washed with RPMI and D-PBS (Fisher Scientific, cat. no. MT21031CV) before staining for cell viability with the Zombie Violet Viability Kit (BioLegend, cat. no. 423113). Next, the appropriate amount of each pMHC complex in a staining buffer of D-PBS with 5% fetal bovine serum (Fisher Scientific, cat. no. MT35016CV) was added to each sample. After 10 min, the aforementioned antibody cocktail was added. After a 30-min incubation period, cells were washed twice in the same staining buffer. All samples were tested in a BD LSRII flow cytometer, and data were analyzed with FlowJo (Treestar). For gating, the lymphocyte population was first selected using forward and side scatter, and then the BV421-negative population was gated out (i.e. excluding dead cells and the CD14, CD19, CD56, and CD4 populations) before selecting the CD3^+^CD8^+^ population. To set for proper gating of dextramer-positive cells, we used cells that were stained with the full antibody panel but no pMHC complexes, and cells that were given the nonhuman pMHC complex.

### TCR sequencing using scRNA-Seq

Cells were stained by following the dextramer procedure with PE-conjugated pMHC complexes only. Cells were sorted by using the BD FACSAria flow cytometer, and PE^+^ cells were collected. V(D)J immune profiling of sorted cells was done with scRNA-Seq, using the 10X Genomics Chromium Single Cell Immune Profiling Workflow at the UCLA Clinical Microarray Core. Each T cell was encapsulated in an oil emulsion droplet with a barcoded gel bead, and reverse transcription was performed to create a barcoded cDNA library. The V(D)J-enriched and gene expression libraries were sequenced using the 10X Genomics Chromium Controller. After sequencing, the Cell Ranger pipeline was used to align reads, filter, count barcodes and assign unique molecular identifiers.

### Next-generation immune repertoire sequencing using the immunoSEQ platform

To assess the T-lymphocyte repertoire of bulk expanded TIL populations, we used the immunoSEQ assay (Adaptive Biotechnologies). This multiplex PCR system uses a mixture of primers that target the rearranged V and J segments of the CDR3 region to assess TCR diversity within a given sample. Genomic DNA from each sample was extracted by using the QIAamp DNA Blood Midi Kit (Qiagen, cat. no. 51185). We provided at least 1 μg of DNA (∼60,000 cells) from each sample to Adaptive Biotechnologies for sequencing at a deep resolution. Resulting sequencing data were analyzed with the immunoSEQ Analyzer Platform (Adaptive Biotechnologies).

### High-throughput αβ TCR pairing using the pairSEQ platform

We provided Adaptive Biotechnologies with frozen bulk expanded TIL samples for their pairSEQ assay, to predict which α and β chains may pair to form a functional TCR. Briefly, T cells were randomly distributed into wells of a 96-well plate. The mRNA was extracted, converted to cDNA, and amplified by using TCR-specific primers. The cDNA of T cells from each well was given a specific barcode, and all wells were pooled together for sequencing. Each TCR sequence was mapped back to the original well through computational demultiplexing. Putative TCR pairs were identified by examining whether a sequenced TCR α chain was frequently seen to share the same well with a specific sequenced TCR β chain, above statistical noise.

## Supporting information

Supplementary Data (flow cytometry)

## Code availability

IRIS source code is accessible on GitHub at https://github.com/Xinglab/IRIS.

## Data availability

The 22 UCLA GBM RNA-Seq data generated for this study were uploaded to BioProject database (BioProject: PRJNA577155). RNA-Seq data used to construct IRIS’s normal and tumor reference panels of AS events are available from the GTEx project (https://gtexportal.org/) and The Cancer Genome Atlas (TCGA) (https://portal.gdc.cancer.gov/legacy-archive/). For the IRIS proteo-transcriptomics analysis, matching RNA-Seq data and MS immunopeptidomics data of B-LCL-S1 and B-LCL-S2 cell lines were retrieved from Laumont *et al.* (GEO: GSM1641206, GSM1641207 and PRIDE: PXD001898). Raw RNA-Seq data of the JeKo-1 lymphoma cell line were obtained from the Cancer Cell Line Encyclopedia via the NCI Genomic Data Commons (https://portal.gdc.cancer.gov/legacy-archive/). Corresponding immunopeptidomics MS data of JeKo-1 were retrieved from Khodadoust *et al.* (PRIDE: PXD004746).

## Acknowledgments

A.R. was funded by the Parker Institute for Cancer Immunotherapy (PICI), the National Institutes of Health (NIH; R35CA197633), and the Ressler Family Fund. O.N.W. was funded by the NIH (R01CA220238, U01CA233074, and P50CA092131) and PICI. Y. Xing was funded by the NIH (R01CA220238 and U01CA233074) and PICI. R.M.P. was funded by the NIH (P50CA211015 and R01CA222695), PICI, and Cancer Research Institute. A.H.L. is a predoctoral fellow supported by the UCLA Tumor Immunology Training Grant (T32CA009120). Flow cytometry and scRNA-Seq experiments were supported in part by funding from the NIH National Center for Advancing Translational Science - UCLA CTSI (UL1TR001881). The authors thank Eric Kutschera (CHOP) for suggestions and assistance with software development, and Life Science Editors for assistance with manuscript editing.

## Author Contributions

Y. Xing conceived the study; Y.P., R.M.P. and Y. Xing designed the research; Y.P. developed the methodology; H.T.Y, Y.W., Y. Xu, J.P., A.C., A.R. and O.N.W. contributed to analytic tools; Y.P., A.H.L., R.M.P. and Y. Xing analyzed data; A.H.L. and C.P. performed experimental validation; and Y.P., A.H.L., K.E.K.-E. and Y. Xing wrote the paper with input from all other authors.

