## Supplementary Figures 1-5 for "IRIS: Big data-informed discovery of cancer immunotherapy targets arising from pre-mRNA alternative splicing"

### Slide 1
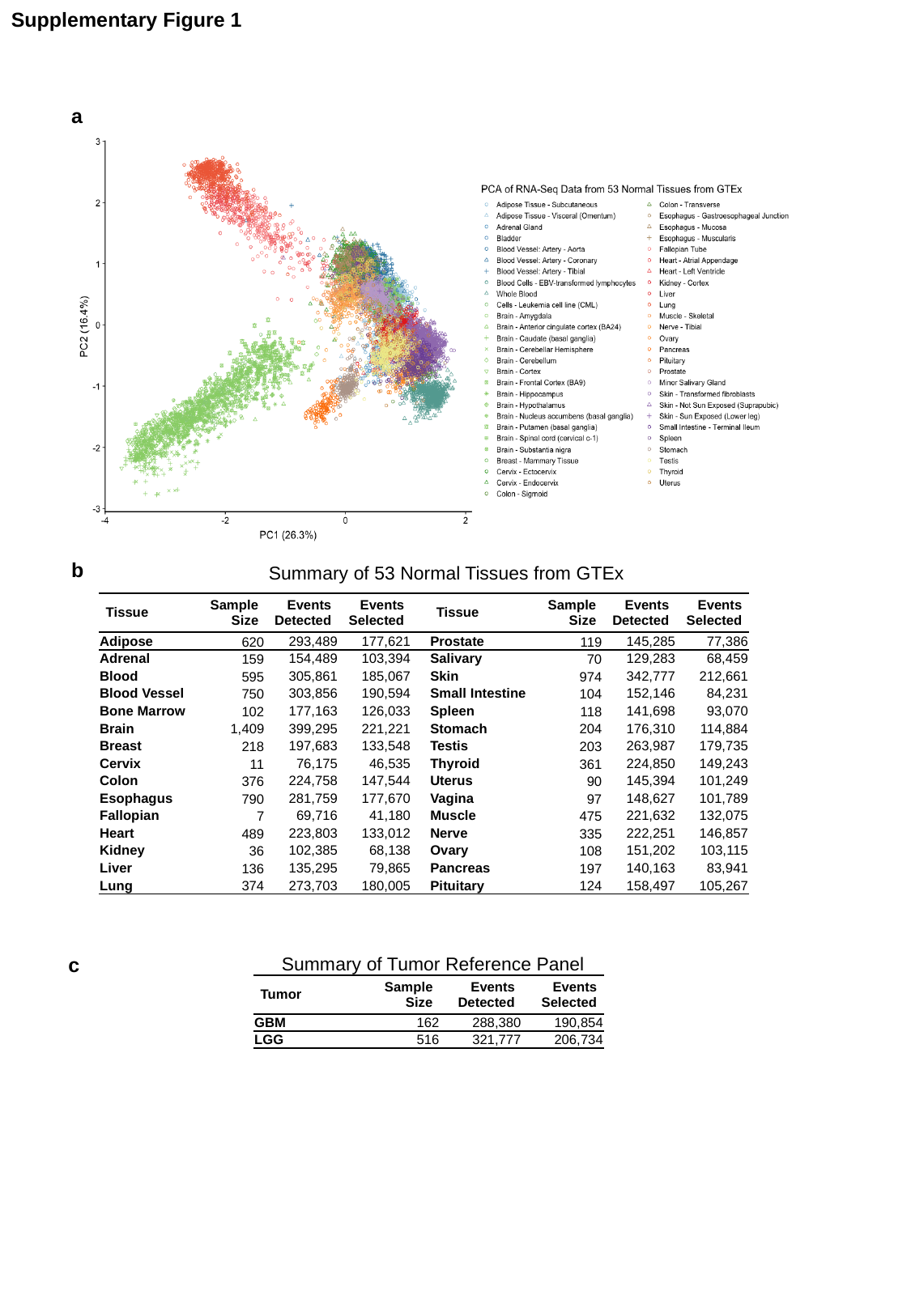

Supplementary Figure 1
a
b
Summary of 53 Normal Tissues from GTEx
| Tissue | Sample Size | Events Detected | Events Selected | | Tissue | Sample Size | Events Detected | Events Selected |
| --- | --- | --- | --- | --- | --- | --- | --- | --- |
| Adipose | 620 | 293,489 | 177,621 | | Prostate | 119 | 145,285 | 77,386 |
| Adrenal | 159 | 154,489 | 103,394 | | Salivary | 70 | 129,283 | 68,459 |
| Blood | 595 | 305,861 | 185,067 | | Skin | 974 | 342,777 | 212,661 |
| Blood Vessel | 750 | 303,856 | 190,594 | | Small Intestine | 104 | 152,146 | 84,231 |
| Bone Marrow | 102 | 177,163 | 126,033 | | Spleen | 118 | 141,698 | 93,070 |
| Brain | 1,409 | 399,295 | 221,221 | | Stomach | 204 | 176,310 | 114,884 |
| Breast | 218 | 197,683 | 133,548 | | Testis | 203 | 263,987 | 179,735 |
| Cervix | 11 | 76,175 | 46,535 | | Thyroid | 361 | 224,850 | 149,243 |
| Colon | 376 | 224,758 | 147,544 | | Uterus | 90 | 145,394 | 101,249 |
| Esophagus | 790 | 281,759 | 177,670 | | Vagina | 97 | 148,627 | 101,789 |
| Fallopian | 7 | 69,716 | 41,180 | | Muscle | 475 | 221,632 | 132,075 |
| Heart | 489 | 223,803 | 133,012 | | Nerve | 335 | 222,251 | 146,857 |
| Kidney | 36 | 102,385 | 68,138 | | Ovary | 108 | 151,202 | 103,115 |
| Liver | 136 | 135,295 | 79,865 | | Pancreas | 197 | 140,163 | 83,941 |
| Lung | 374 | 273,703 | 180,005 | | Pituitary | 124 | 158,497 | 105,267 |
c
Summary of Tumor Reference Panel
| Tumor | Sample Size | Events Detected | Events Selected |
| --- | --- | --- | --- |
| GBM | 162 | 288,380 | 190,854 |
| LGG | 516 | 321,777 | 206,734 |

### Slide 2
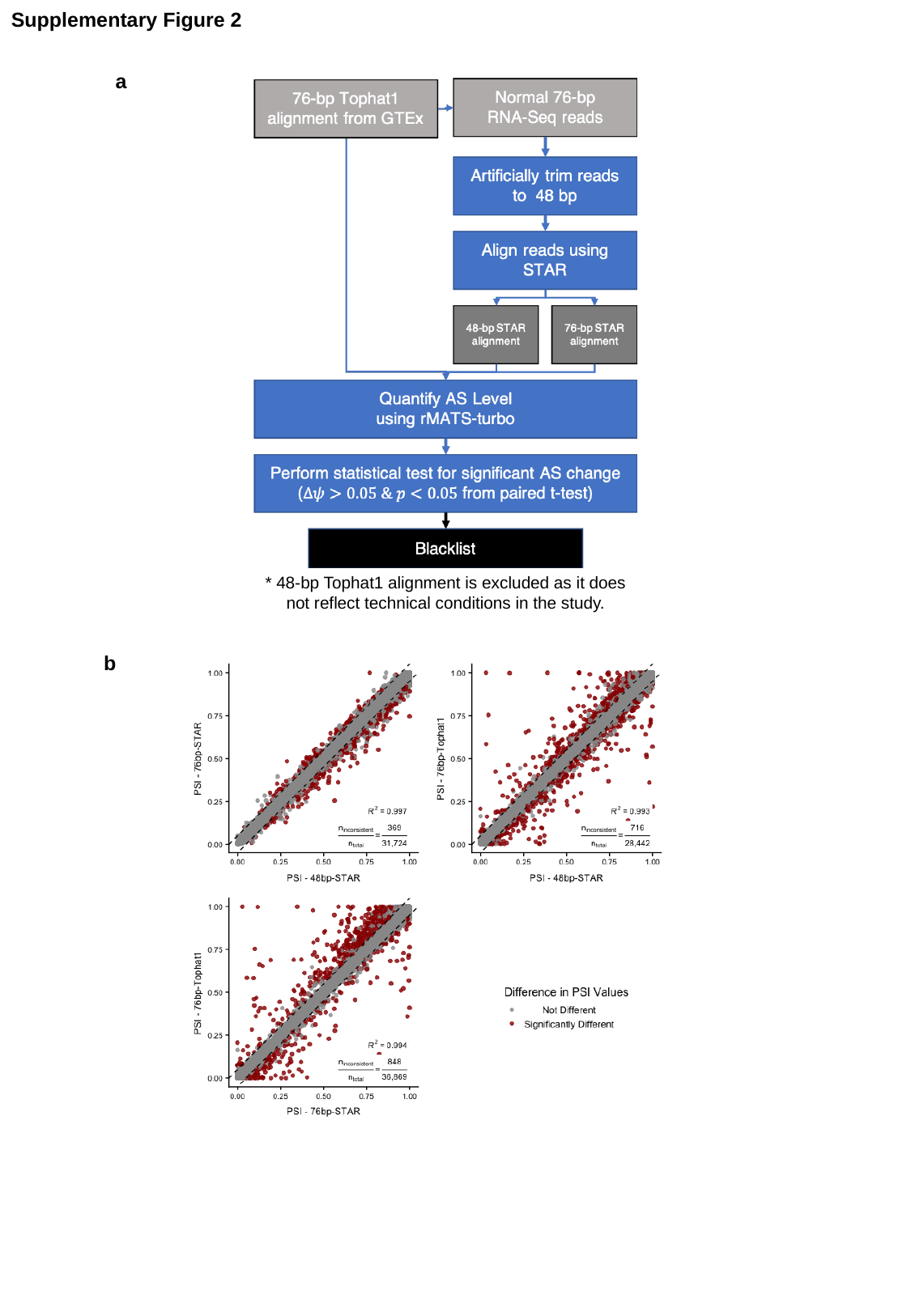

Supplementary Figure 2
a
* 48-bp Tophat1 alignment is excluded as it does not reflect technical conditions in the study.
b

### Slide 3
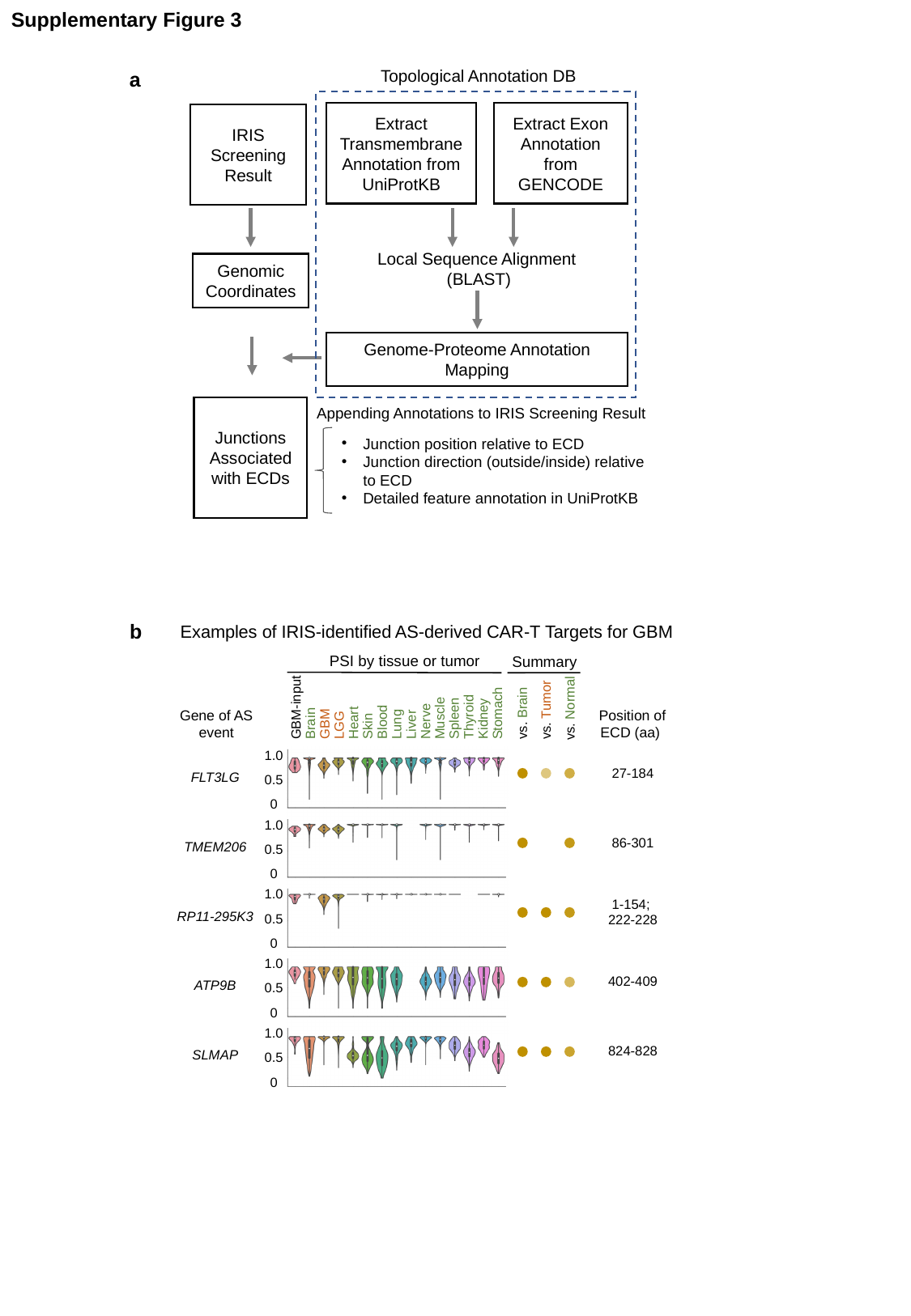

Supplementary Figure 3
Topological Annotation DB
Extract Exon Annotation from GENCODE
Extract Transmembrane Annotation from UniProtKB
IRIS Screening
Result
Local Sequence Alignment
(BLAST)
Genomic Coordinates
Genome-Proteome Annotation Mapping
Junctions Associated with ECDs
Appending Annotations to IRIS Screening Result
Junction position relative to ECD
Junction direction (outside/inside) relative to ECD
Detailed feature annotation in UniProtKB
a
b
Examples of IRIS-identified AS-derived CAR-T Targets for GBM
PSI by tissue or tumor
GBM-input
Stomach
Thyroid
Muscle
Spleen
Kidney
Nerve
Blood
Heart
Brain
GBM
Lung
Liver
LGG
Skin
vs. Normal
vs. Tumor
vs. Brain
1.0
0.5
0
1.0
0.5
0
1.0
0.5
0
1.0
0.5
0
1.0
0.5
0
Summary
Gene of AS event
Position of ECD (aa)
| 27-184 |
| --- |
| 86-301 |
| 1-154; 222-228 |
| 402-409 |
| 824-828 |
| FLT3LG |
| --- |
| TMEM206 |
| RP11-295K3 |
| ATP9B |
| SLMAP |

### Slide 4
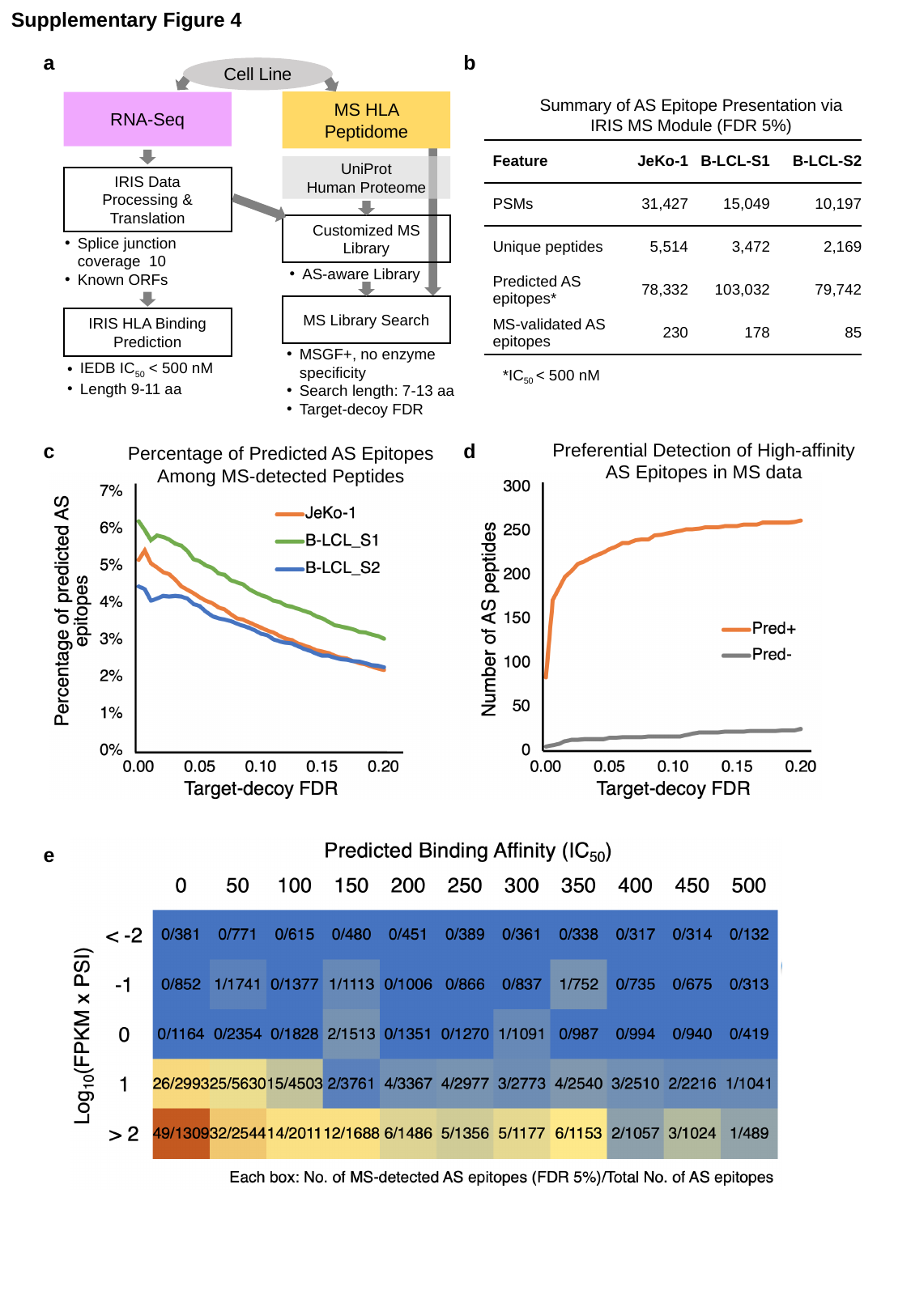

Supplementary Figure 4
a
b
Cell Line
MS HLA Peptidome
RNA-Seq
UniProt
Human Proteome
IRIS Data Processing & Translation
Customized MS Library
AS-aware Library
MS Library Search
IRIS HLA Binding Prediction
IEDB IC_50_ < 500 nM
Length 9-11 aa
MSGF+, no enzyme specificity
Search length: 7-13 aa
Target-decoy FDR
Summary of AS Epitope Presentation via
IRIS MS Module (FDR 5%)
| Feature | JeKo-1 | B-LCL-S1 | B-LCL-S2 |
| --- | --- | --- | --- |
| PSMs | 31,427 | 15,049 | 10,197 |
| Unique peptides | 5,514 | 3,472 | 2,169 |
| Predicted AS epitopes\* | 78,332 | 103,032 | 79,742 |
| MS-validated AS epitopes | 230 | 178 | 85 |
*IC_50_ < 500 nM
Preferential Detection of High-affinity AS Epitopes in MS data
c
d
Percentage of Predicted AS Epitopes Among MS-detected Peptides
e

### Slide 5
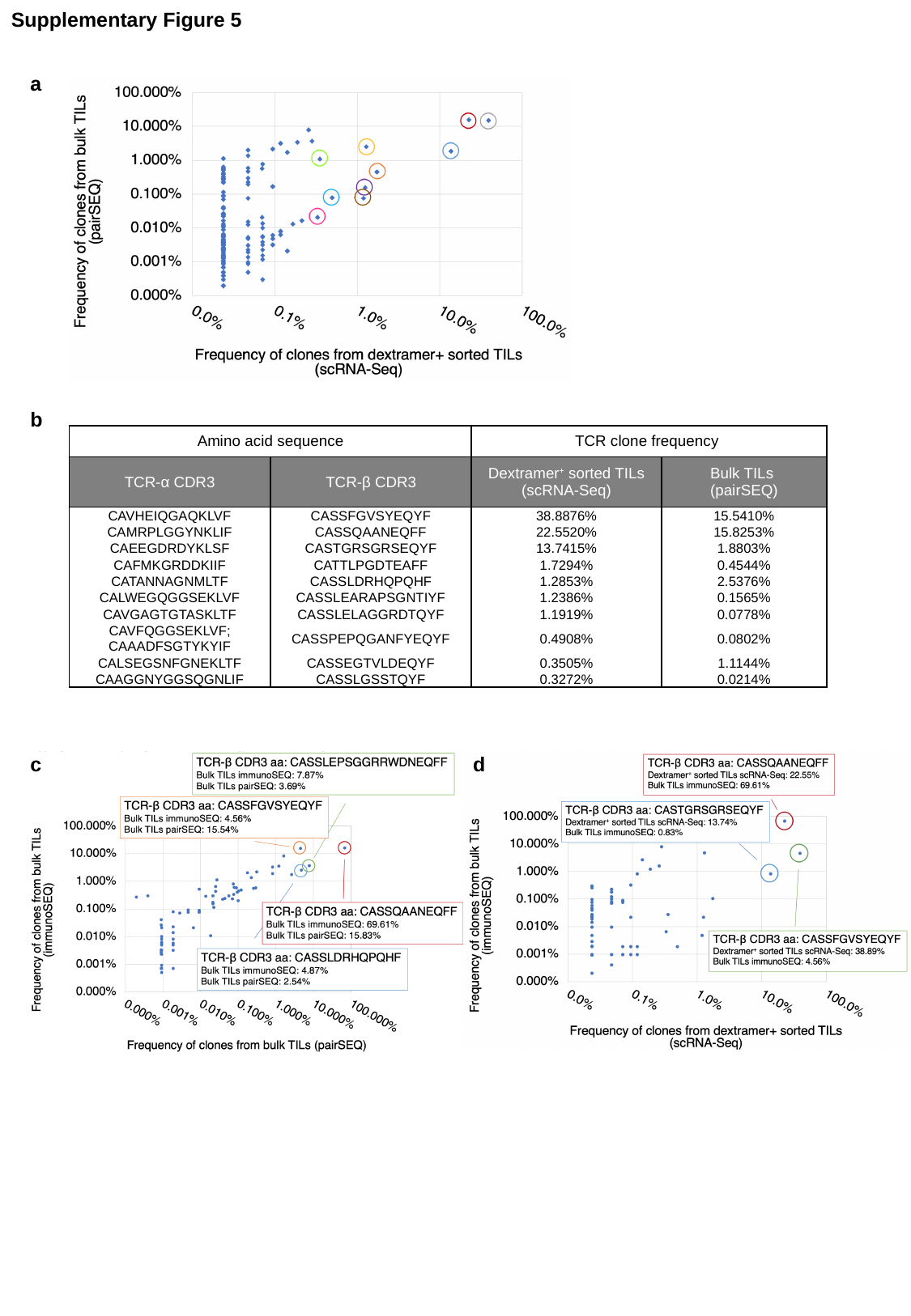

Supplementary Figure 5
a
b
| Amino acid sequence | | TCR clone frequency | |
| --- | --- | --- | --- |
| TCR-α CDR3 | TCR-β CDR3 | Dextramer^+^ sorted TILs (scRNA-Seq) | Bulk TILs (pairSEQ) |
| CAVHEIQGAQKLVF | CASSFGVSYEQYF | 38.8876% | 15.5410% |
| CAMRPLGGYNKLIF | CASSQAANEQFF | 22.5520% | 15.8253% |
| CAEEGDRDYKLSF | CASTGRSGRSEQYF | 13.7415% | 1.8803% |
| CAFMKGRDDKIIF | CATTLPGDTEAFF | 1.7294% | 0.4544% |
| CATANNAGNMLTF | CASSLDRHQPQHF | 1.2853% | 2.5376% |
| CALWEGQGGSEKLVF | CASSLEARAPSGNTIYF | 1.2386% | 0.1565% |
| CAVGAGTGTASKLTF | CASSLELAGGRDTQYF | 1.1919% | 0.0778% |
| CAVFQGGSEKLVF; CAAADFSGTYKYIF | CASSPEPQGANFYEQYF | 0.4908% | 0.0802% |
| CALSEGSNFGNEKLTF | CASSEGTVLDEQYF | 0.3505% | 1.1144% |
| CAAGGNYGGSQGNLIF | CASSLGSSTQYF | 0.3272% | 0.0214% |
c
d
