## Supplementary Figure Legends for "IRIS: Big data-informed discovery of cancer immunotherapy targets arising from pre-mRNA alternative splicing"

**Supplementary Figure 1. RNA-Seq big-data reference panels in IRIS.**

**a,** Exon-based principal component analysis (PCA) of RNA-Seq data of 9,662 samples from 53 normal tissues from the GTEx consortium. Samples from the same histological site are grouped by color. Samples from different subregions of the same histological site are differentiated by different shapes.

**b,** Summary of 53 normal tissues from the GTEx consortium. Data for all 53 tissues are available to IRIS users as a reference panel of normal tissues. In the present study, 11 selected vital tissues (heart, skin, blood, lung, liver, nerve, muscle, spleen, thyroid, kidney, and stomach) were used for the ‘normal panel’. ‘Events Selected’ represent AS events with an average count ≥ 10 reads for the sum of all splice junctions across all samples in that tissue.

**c,** Summary of the tumor reference panel (TCGA tumor samples relevant to GBM). ‘Events Selected’ represent AS events with an average count ≥ 10 reads for the sum of all splice junctions across all samples in that tumor type.

**Supplementary Figure 2. Identification of AS events that are prone to measurement errors due to technical variances across big-data reference panels.**

**a,** Computational workflow to create a ‘blacklist’ of error-prone AS events. Normal 76-bp RNA-Seq reads were artificially trimmed to 48 bp. RNA-Seq files (76- and 48-bp) were aligned by using two different aligners (Tophat and STAR). AS events were quantified by rMATS-turbo. AS events with statistically significant differences in PSI values among RNA-Seq datasets with distinct technical conditions were identified and included in a blacklist.

**b,** Scatter plots comparing PSI values of GTEx normal brain RNA-Seq data estimated under distinct technical conditions (read lengths: 48- and 76-bp, aligners: STAR and Tophat). ‘Significantly different’ AS events were defined as those with significantly different PSI values (p < 0.05, abs(Δψ) > 0.05 from paired *t*-test).

**Supplementary Figure 3. CAR-T target prediction by IRIS.**

**a,** Computational workflow to annotate protein extracellular domain (ECD)-associated AS events for CAR-T target discovery.

**b,** Five examples of IRIS-identified AS-derived CAR-T targets for 22 GBM samples. Position of the ECD in amino acid (aa) sequence was obtained from UniProtKB.

**Supplementary Figure 4. Proteo-transcriptomic analysis of HLA presentation of AS-derived epitopes in normal and tumor cell lines.**

**a,** Proteo-transcriptomics workflow adopted by IRIS to discover splice-junction peptides in MS datasets. IRIS inputs MS data (right), such as whole-cell proteomics, surfaceomics, or immunopeptidomics (HLA peptidomics) data. RNA-Seq-based custom proteome library is constructed and searched using MSGF+.

**b,** Summary of HLA presentation of AS-derived epitopes in JeKo-1 (lymphoma) and B-LCL (normal) cell lines. Peptide-spectrum matches (‘PSMs’) and ‘Unique peptides’ are provided by MSGF+ with a target-decoy FDR of 5%. ‘Predicted AS epitopes’ are generated by the IRIS prediction module, which utilizes IEDB predictors. AS epitopes that are predicted by IRIS and detected in the MS data are considered ‘MS-validated AS epitopes’.

**c,** Percentage of IRIS-predicted AS-derived epitopes among all MS-detected peptides. Graph shows the percentage of all MS-detected peptides that are IRIS-predicted AS-derived epitopes (y-axis) as a function of the MSGF+ target-decoy FDR (x-axis).

**d,** Preferential detection of high-affinity AS-derived peptides in MS data. Graph shows the number of AS-derived peptides detected in JeKo-1 MS data (y-axis) as a function of the MSGF+ target-decoy FDR (x-axis). Peptides with high (IC_50_ < 500 nM; Pred+, orange) and low (IC_50_ > 500 nM; Pred-, grey) predicted HLA binding affinities are shown.

**e,** Heatmap depiction of distribution of AS-derived epitopes in JeKo-1 MS immunopeptidome, as a function of predicted HLA binding affinity and transcript expression level. AS-derived peptides are binned by the corresponding transcripts’ expression levels and IEDB-predicted binding affinity scores. Heatmap is colored from red (high) to yellow (90^th^ percentile) to blue (low), reflecting the proportion of IRIS-predicted AS-derived epitopes that are MS-detected in each bin.

**Supplementary Figure 5. Consistent distributions of high-frequency TCR clones in one patient’s TIL population revealed by multiple TCR sequencing approaches.**

**a,** Scatter plot comparing scRNA-Seq and bulk TIL pairSEQ for detection of high-frequency TCR clones. Graph shows frequency detected from bulk TIL samples using pairSEQ (y-axis) and scRNA-Seq on dextramer-positive sorted TIL samples (x-axis). As a complementary validation of scRNA-Seq, clonotypes from pairSEQ were matched to scRNA-Seq results by either CDR3 pairs or β chains, whichever matched best. The 10 most abundant TCR clones by scRNA-Seq that overlapped with clones detected by bulk TIL pairSEQ are circled.

**b,** Table showing CDR3 amino acid sequences of the 10 most abundant TCR clones detected by scRNA-Seq and their corresponding detection frequencies by bulk TIL pairSEQ. As a complementary validation of scRNA-Seq, clonotypes from pairSEQ were matched to scRNA-Seq results by either CDR3 pairs or β chains, whichever matched best.

**c,** Scatter plot comparing bulk TIL immunoSEQ and bulk TIL pairSEQ for detection of high-frequency TCR clones. Graph shows frequency detected from bulk TIL samples using immunoSEQ (y-axis) and pairSEQ (x-axis). Clonotypes from immunoSEQ were matched to pairSEQ results by the best CDR3 β chains. Four high-frequency overlapping clones from both methods are circled and color-coded, with β-chain CDR3 amino acid sequences and frequencies by each method shown in boxes.

**d,** Scatter plot comparing scRNA-Seq and bulk TIL immunoSEQ for detection of high-frequency TCR clones. Graph shows frequency detected from bulk TIL samples using immunoSEQ (y-axis) and scRNA-Seq on dextramer-positive sorted TIL samples (x-axis). As a complementary validation of scRNA-Seq, clonotypes from immunoSEQ were matched to scRNA-Seq results by the best CDR3 β chains. Three high-frequency overlapping clones from both methods are circled and color-coded, with β-chain CDR3 amino acid sequences and frequencies by each method shown in boxes.
