## Supplementary Data (flow cytometry) for "IRIS: Big data-informed discovery of cancer immunotherapy targets arising from pre-mRNA alternative splicing"

### Slide 1
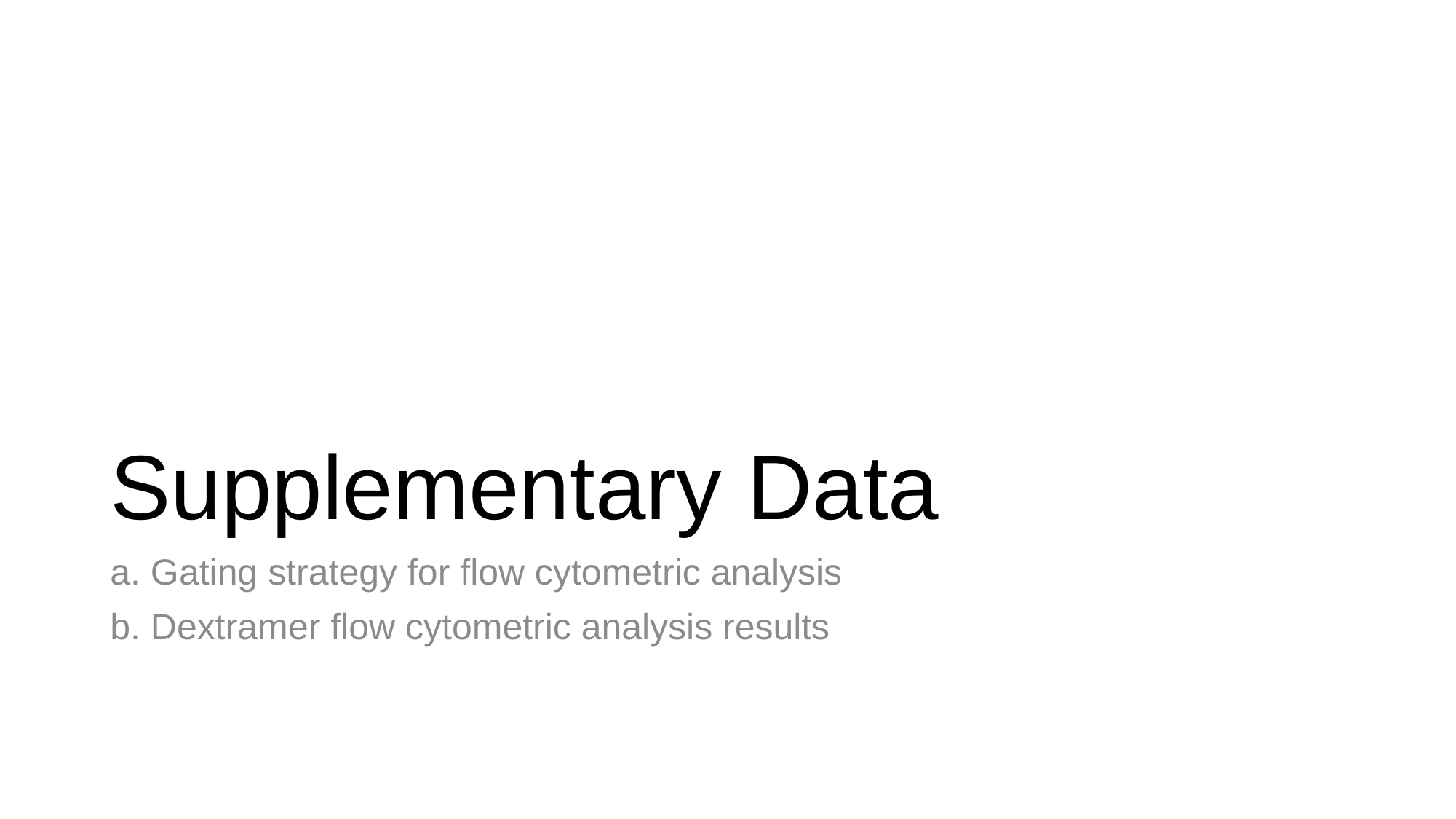

#### Supplementary Data
a. Gating strategy for flow cytometric analysis
b. Dextramer flow cytometric analysis results

### Slide 2
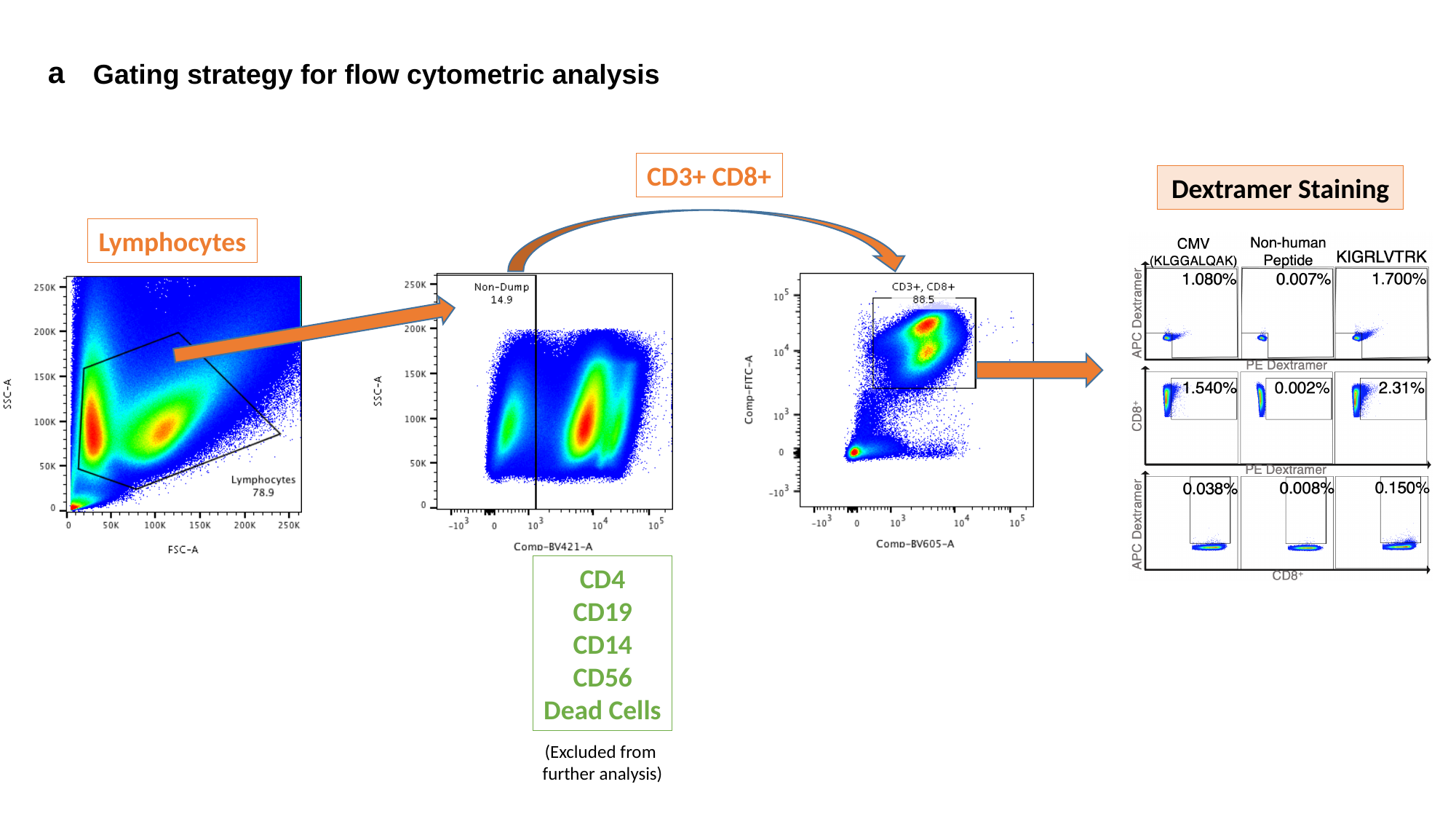

a
Gating strategy for flow cytometric analysis
CD3+ CD8+
Dextramer Staining
Lymphocytes
CD4
CD19
CD14
CD56
Dead Cells
(Excluded from
further analysis)

### Slide 3
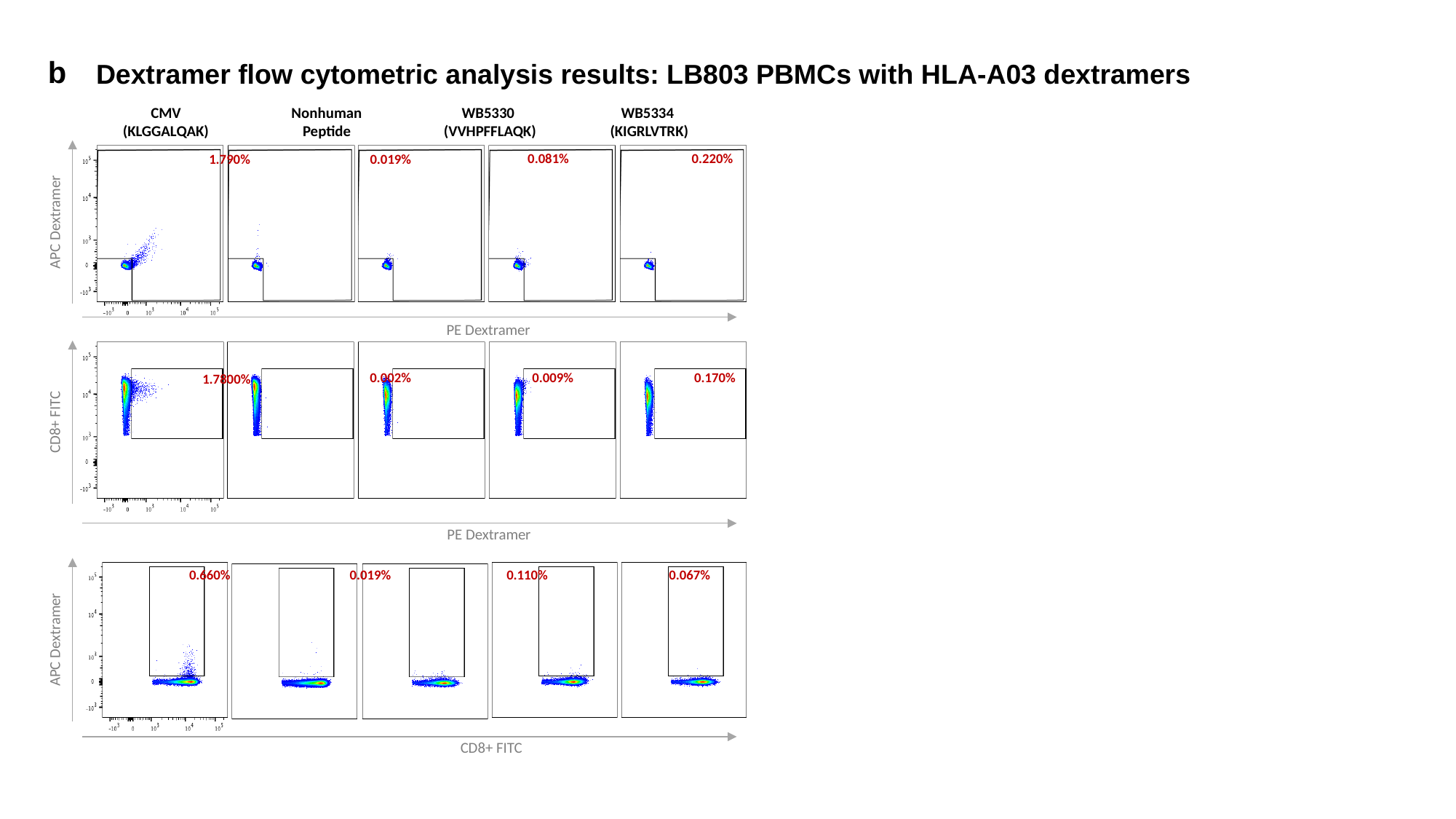

b
Dextramer flow cytometric analysis results: LB803 PBMCs with HLA-A03 dextramers
CMV
(KLGGALQAK)
Nonhuman
Peptide
WB5330
(VVHPFFLAQK)
WB5334
(KIGRLVTRK)
0.220%
0.081%
0.019%
1.790%
APC Dextramer
PE Dextramer
0.002%
0.009%
0.170%
1.7800%
CD8+ FITC
PE Dextramer
0.660%
0.019%
0.110%
0.067%
APC Dextramer
CD8+ FITC

### Slide 4
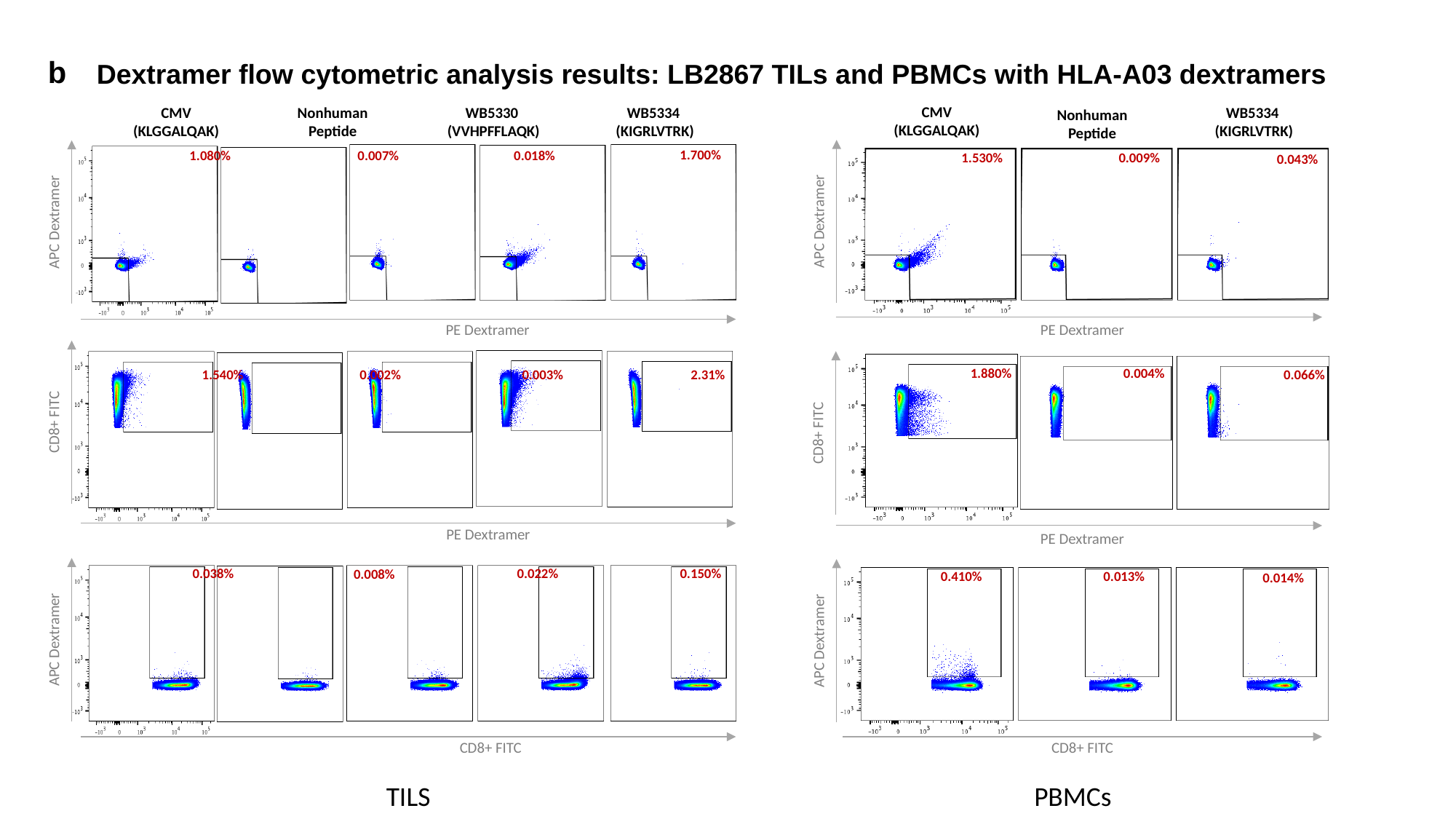

b
Dextramer flow cytometric analysis results: LB2867 TILs and PBMCs with HLA-A03 dextramers
CMV
(KLGGALQAK)
WB5334
(KIGRLVTRK)
Nonhuman
Peptide
1.530%
0.009%
0.043%
APC Dextramer
PE Dextramer
1.880%
0.004%
0.066%
CD8+ FITC
PE Dextramer
0.410%
0.013%
0.014%
APC Dextramer
CD8+ FITC
CMV
(KLGGALQAK)
Nonhuman
Peptide
WB5330
(VVHPFFLAQK)
WB5334
(KIGRLVTRK)
1.700%
0.018%
0.007%
1.080%
APC Dextramer
PE Dextramer
1.540%
0.002%
0.003%
2.31%
CD8+ FITC
PE Dextramer
0.038%
0.022%
0.150%
0.008%
APC Dextramer
CD8+ FITC
PBMCs
TILS

### Slide 5
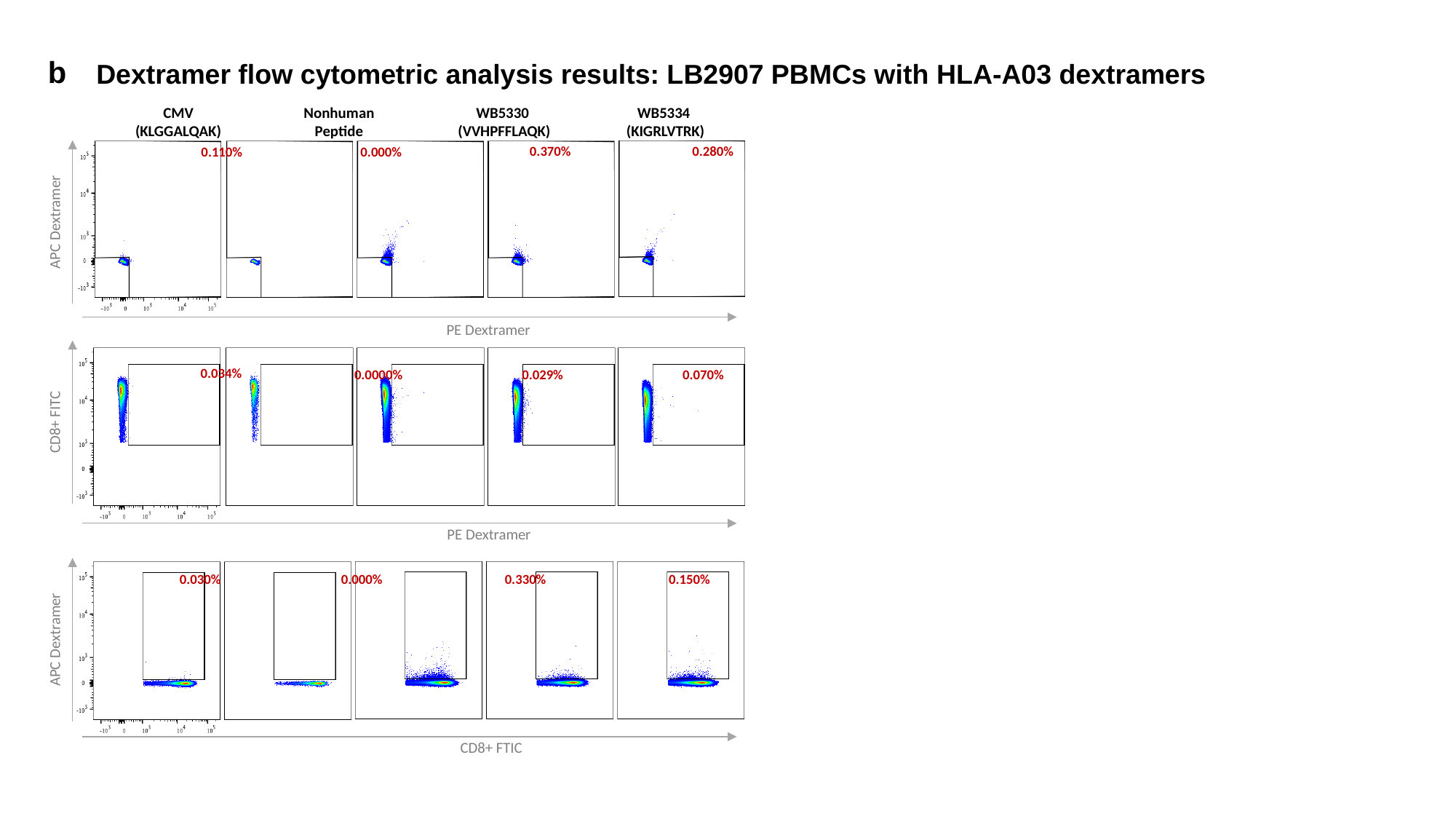

b
Dextramer flow cytometric analysis results: LB2907 PBMCs with HLA-A03 dextramers
CMV
(KLGGALQAK)
Nonhuman
Peptide
WB5334
(KIGRLVTRK)
WB5330
(VVHPFFLAQK)
0.280%
0.370%
0.000%
0.110%
APC Dextramer
PE Dextramer
0.034%
0.0000%
0.029%
0.070%
CD8+ FITC
PE Dextramer
0.030%
0.330%
0.150%
0.000%
APC Dextramer
CD8+ FTIC

### Slide 6
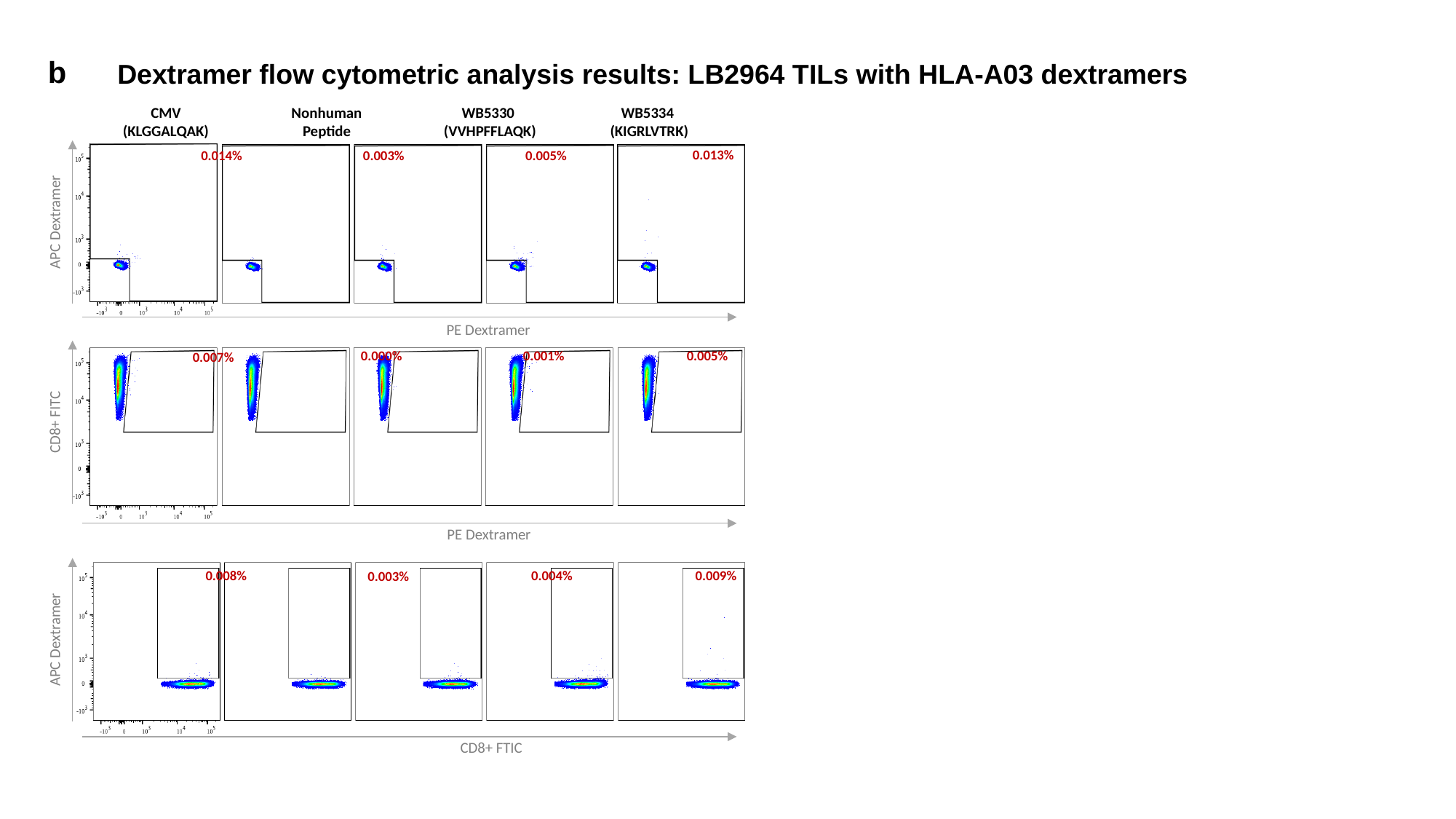

b
Dextramer flow cytometric analysis results: LB2964 TILs with HLA-A03 dextramers
CMV
(KLGGALQAK)
Nonhuman
Peptide
WB5330
(VVHPFFLAQK)
WB5334
(KIGRLVTRK)
0.013%
0.005%
0.003%
0.014%
APC Dextramer
PE Dextramer
0.000%
0.001%
0.005%
0.007%
CD8+ FITC
PE Dextramer
0.008%
0.004%
0.009%
0.003%
APC Dextramer
CD8+ FTIC

### Slide 7
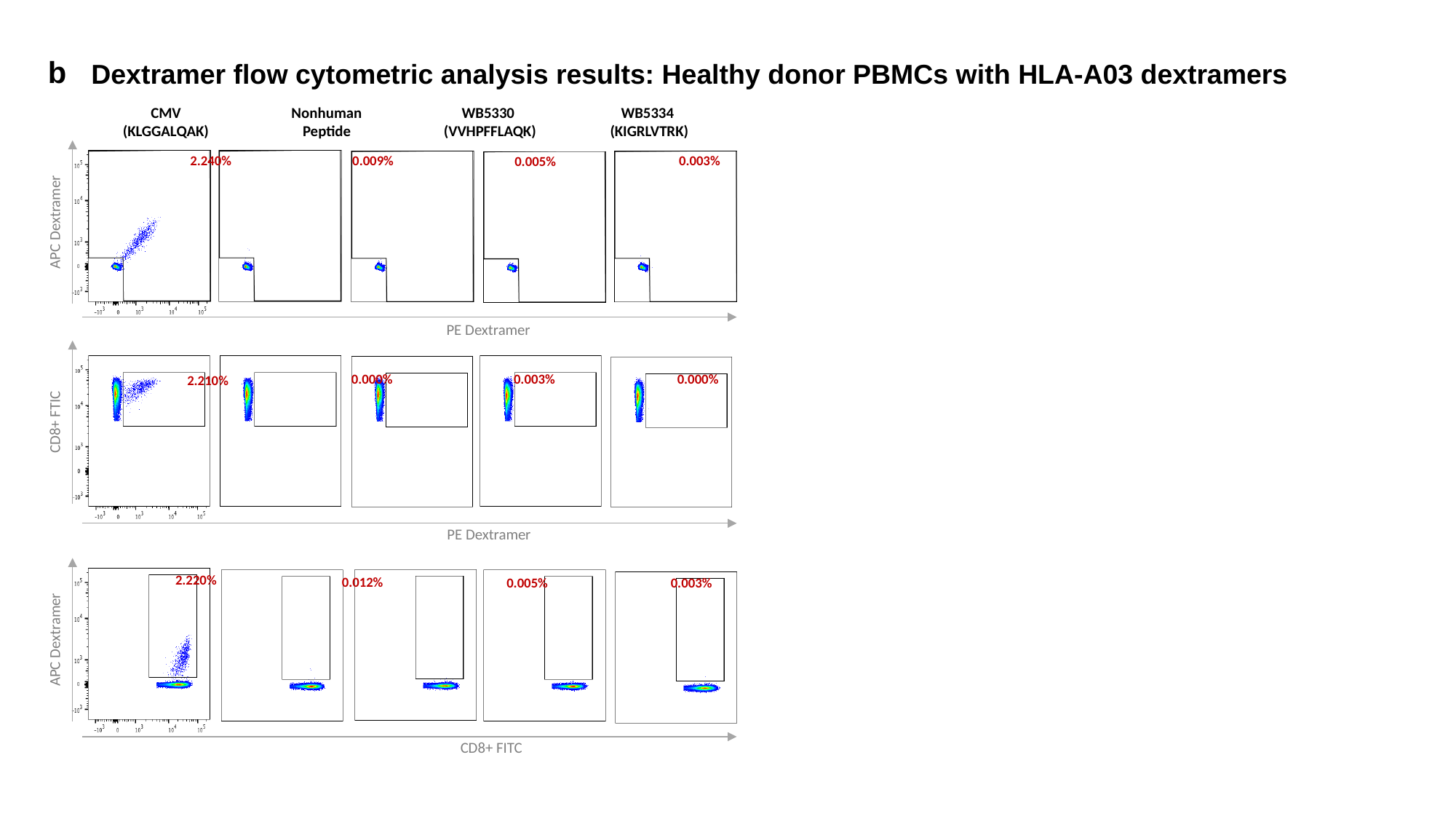

b
Dextramer flow cytometric analysis results: Healthy donor PBMCs with HLA-A03 dextramers
CMV
(KLGGALQAK)
Nonhuman
Peptide
WB5330
(VVHPFFLAQK)
WB5334
(KIGRLVTRK)
0.009%
0.003%
2.240%
0.005%
APC Dextramer
PE Dextramer
0.000%
0.003%
0.000%
2.210%
CD8+ FTIC
PE Dextramer
2.220%
0.012%
0.005%
0.003%
APC Dextramer
CD8+ FITC

### Slide 8
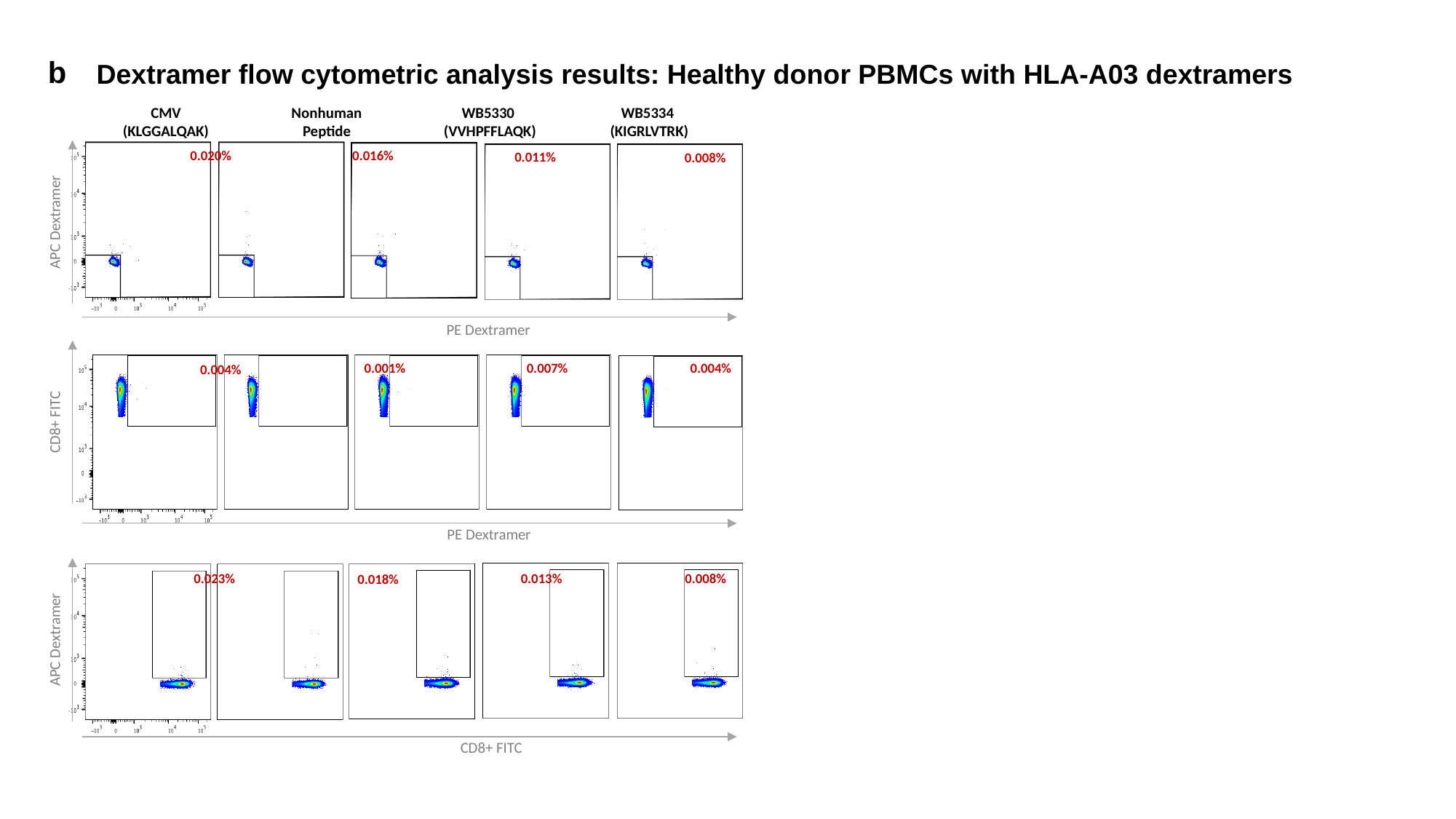

b
Dextramer flow cytometric analysis results: Healthy donor PBMCs with HLA-A03 dextramers
CMV
(KLGGALQAK)
Nonhuman
Peptide
WB5330
(VVHPFFLAQK)
WB5334
(KIGRLVTRK)
0.016%
0.020%
0.011%
0.008%
APC Dextramer
PE Dextramer
0.001%
0.007%
0.004%
0.004%
CD8+ FITC
PE Dextramer
0.023%
0.013%
0.008%
0.018%
APC Dextramer
CD8+ FITC

### Slide 9
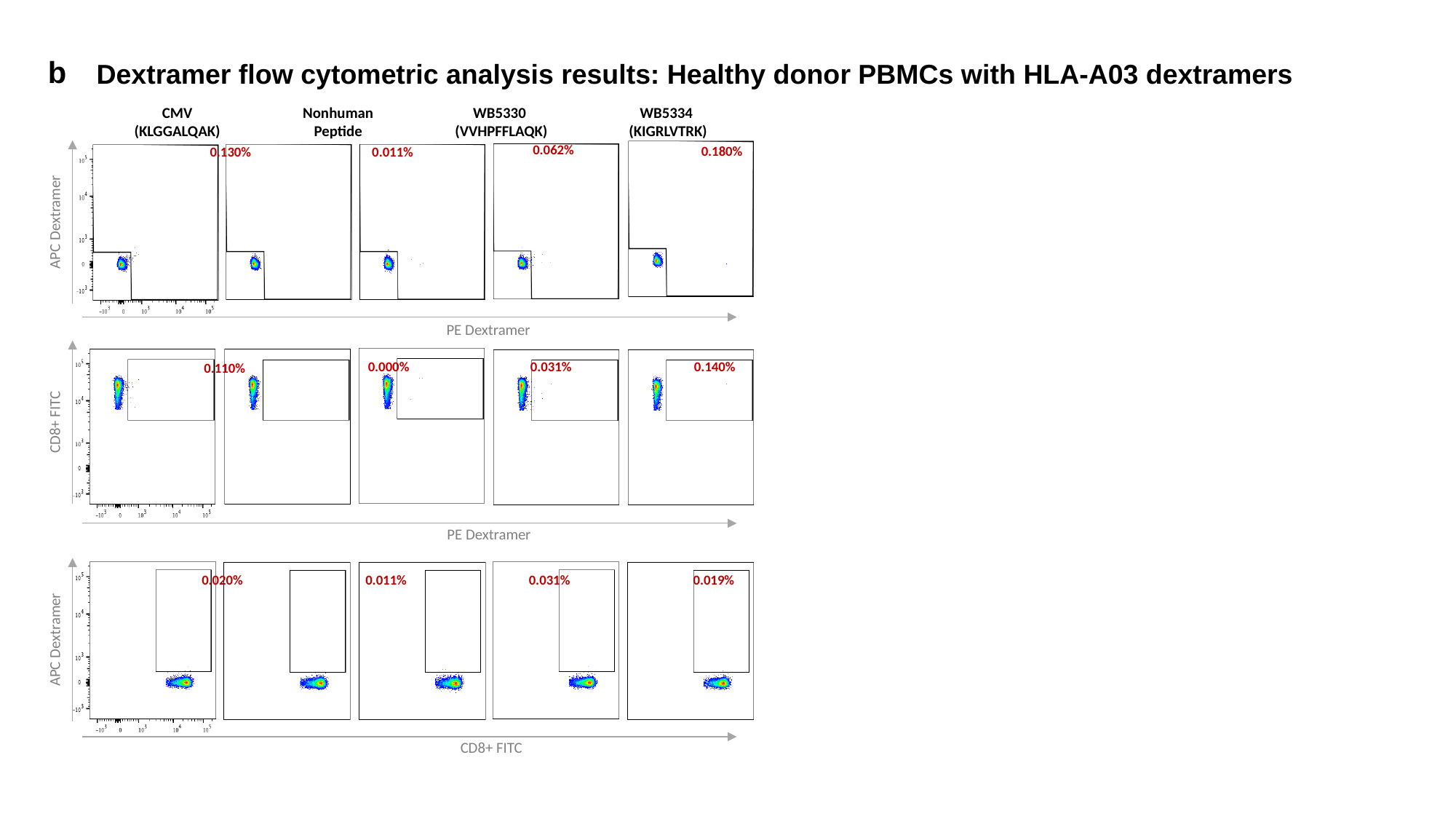

b
Dextramer flow cytometric analysis results: Healthy donor PBMCs with HLA-A03 dextramers
CMV
(KLGGALQAK)
Nonhuman
Peptide
WB5330
(VVHPFFLAQK)
WB5334
(KIGRLVTRK)
0.062%
0.180%
0.011%
0.130%
APC Dextramer
PE Dextramer
0.000%
0.031%
0.140%
0.110%
CD8+ FITC
PE Dextramer
0.020%
0.031%
0.019%
0.011%
APC Dextramer
CD8+ FITC

### Slide 10
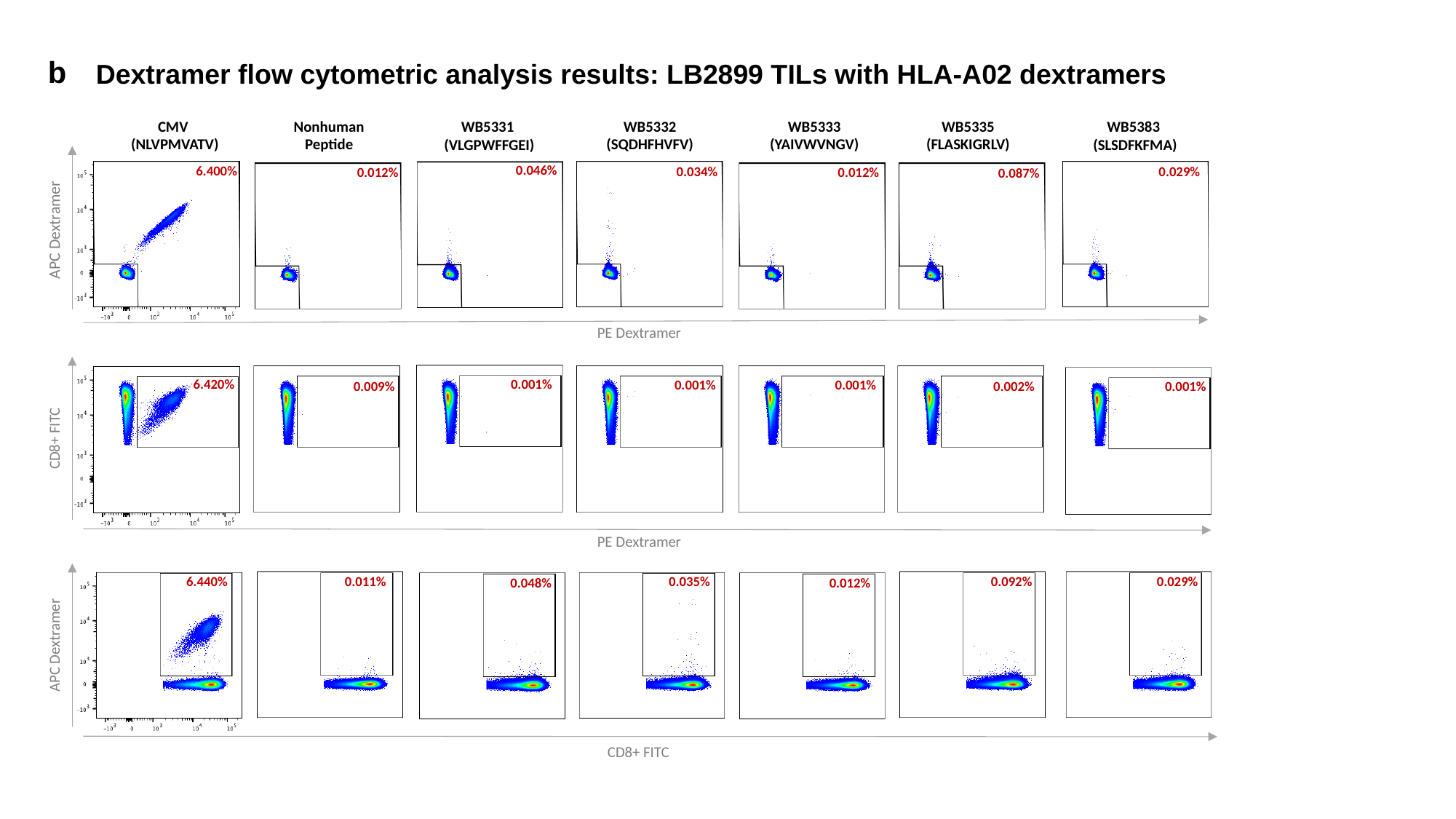

b
Dextramer flow cytometric analysis results: LB2899 TILs with HLA-A02 dextramers
CMV
(NLVPMVATV)
Nonhuman
Peptide
WB5332
(SQDHFHVFV)
WB5333
(YAIVWVNGV)
WB5335
(FLASKIGRLV)
WB5383
(SLSDFKFMA)
WB5331
(VLGPWFFGEI)
0.046%
6.400%
0.034%
0.029%
0.012%
0.012%
0.087%
APC Dextramer
PE Dextramer
6.420%
0.001%
0.001%
0.001%
0.002%
0.001%
0.009%
CD8+ FITC
PE Dextramer
0.035%
0.092%
0.029%
6.440%
0.011%
0.012%
0.048%
APC Dextramer
CD8+ FITC

### Slide 11
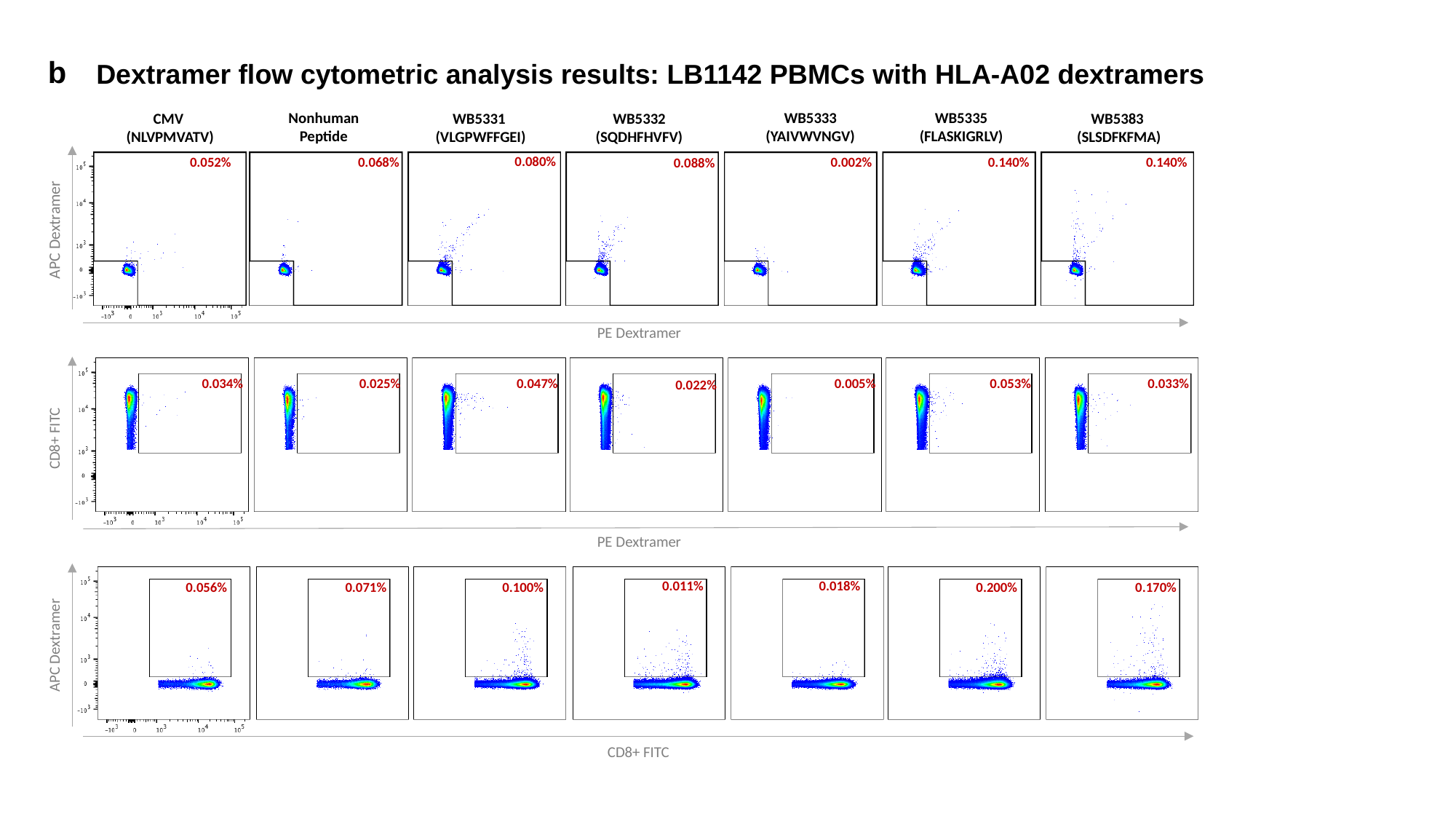

b
Dextramer flow cytometric analysis results: LB1142 PBMCs with HLA-A02 dextramers
Nonhuman
Peptide
WB5333
(YAIVWVNGV)
WB5335
(FLASKIGRLV)
WB5383
(SLSDFKFMA)
WB5332
(SQDHFHVFV)
CMV
(NLVPMVATV)
WB5331
(VLGPWFFGEI)
0.080%
0.052%
0.068%
0.002%
0.140%
0.140%
0.088%
APC Dextramer
PE Dextramer
0.047%
0.034%
0.025%
0.005%
0.033%
0.053%
0.022%
CD8+ FITC
PE Dextramer
0.011%
0.018%
0.056%
0.071%
0.200%
0.170%
0.100%
APC Dextramer
CD8+ FITC

### Slide 12
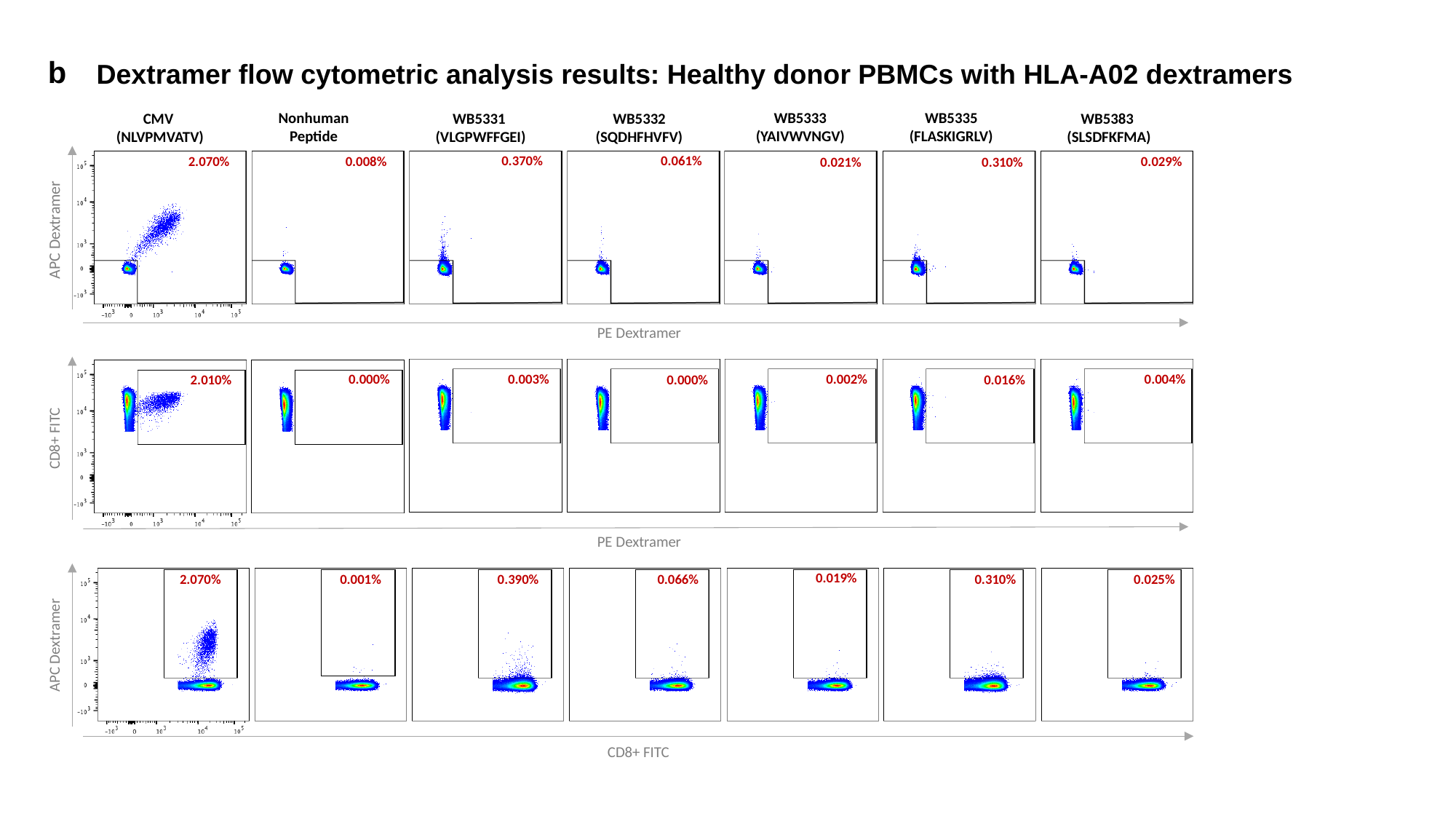

b
Dextramer flow cytometric analysis results: Healthy donor PBMCs with HLA-A02 dextramers
Nonhuman
Peptide
WB5333
(YAIVWVNGV)
WB5335
(FLASKIGRLV)
WB5383
(SLSDFKFMA)
WB5332
(SQDHFHVFV)
CMV
(NLVPMVATV)
WB5331
(VLGPWFFGEI)
0.061%
0.370%
0.029%
2.070%
0.008%
0.021%
0.310%
APC Dextramer
PE Dextramer
0.002%
0.004%
0.003%
0.000%
0.000%
0.016%
2.010%
CD8+ FITC
PE Dextramer
0.019%
0.066%
0.310%
0.025%
2.070%
0.001%
0.390%
APC Dextramer
CD8+ FITC

### Slide 13
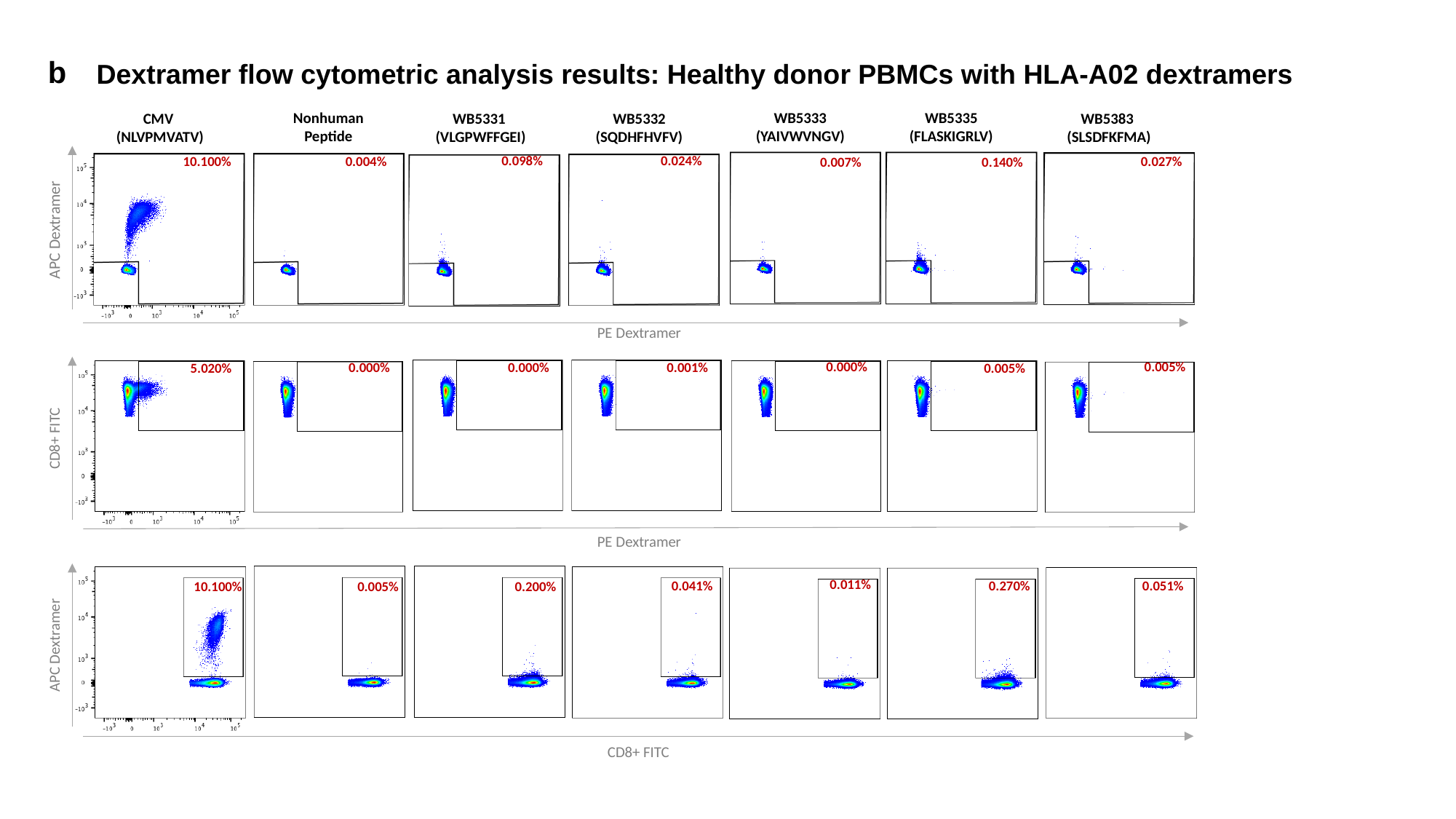

b
Dextramer flow cytometric analysis results: Healthy donor PBMCs with HLA-A02 dextramers
Nonhuman
Peptide
WB5333
(YAIVWVNGV)
WB5335
(FLASKIGRLV)
WB5383
(SLSDFKFMA)
WB5332
(SQDHFHVFV)
CMV
(NLVPMVATV)
WB5331
(VLGPWFFGEI)
0.024%
0.098%
0.027%
10.100%
0.004%
0.007%
0.140%
APC Dextramer
PE Dextramer
0.000%
0.005%
0.000%
0.000%
0.001%
0.005%
5.020%
CD8+ FITC
PE Dextramer
0.011%
0.041%
0.270%
0.051%
10.100%
0.005%
0.200%
APC Dextramer
CD8+ FITC

### Slide 14
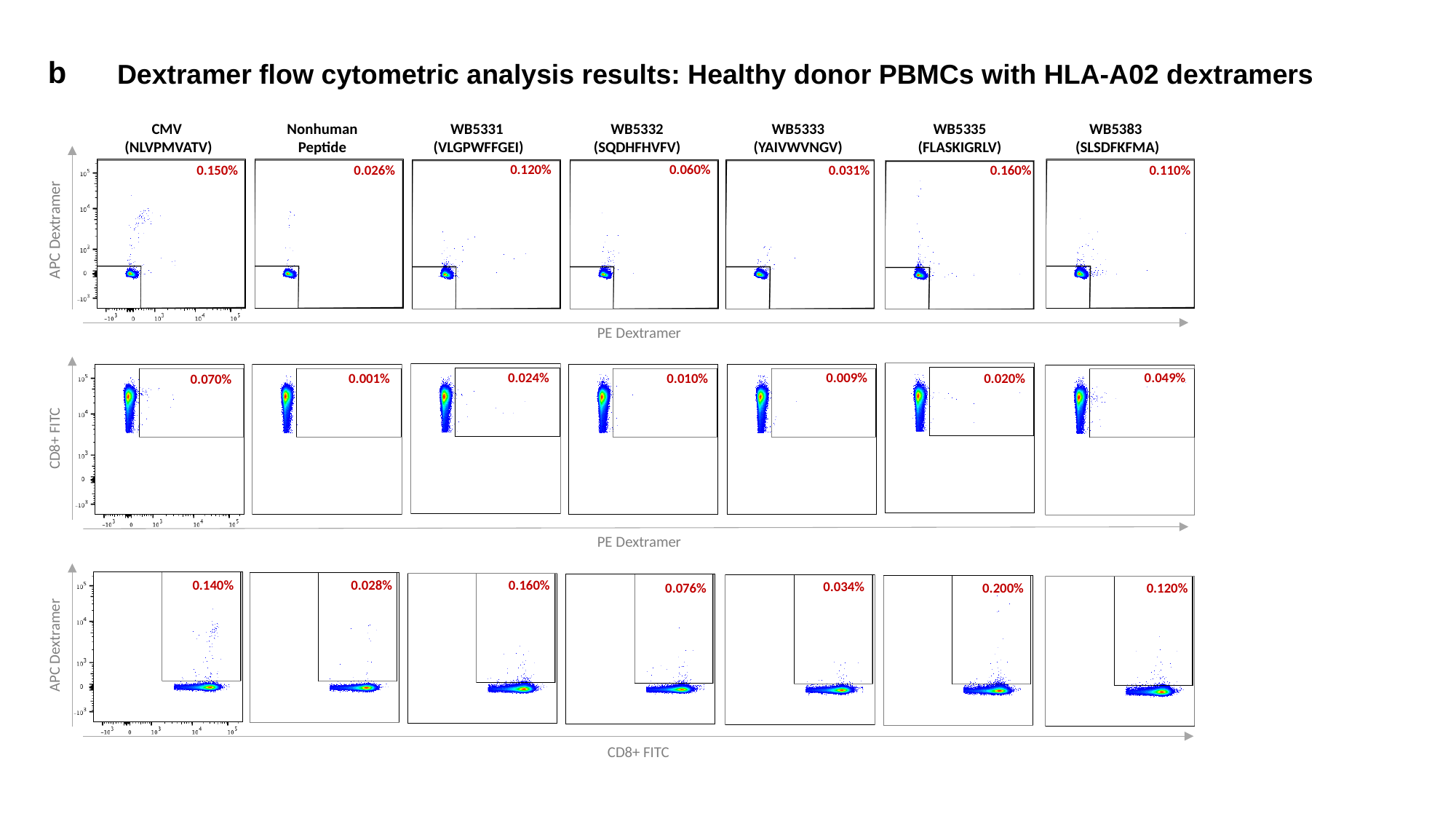

b
Dextramer flow cytometric analysis results: Healthy donor PBMCs with HLA-A02 dextramers
Nonhuman
Peptide
WB5333
(YAIVWVNGV)
WB5335
(FLASKIGRLV)
WB5383
(SLSDFKFMA)
WB5332
(SQDHFHVFV)
CMV
(NLVPMVATV)
WB5331
(VLGPWFFGEI)
0.060%
0.120%
0.110%
0.150%
0.026%
0.031%
0.160%
APC Dextramer
PE Dextramer
0.009%
0.049%
0.024%
0.001%
0.010%
0.020%
0.070%
CD8+ FITC
PE Dextramer
0.140%
0.028%
0.160%
0.034%
0.076%
0.200%
0.120%
APC Dextramer
CD8+ FITC
